# Extracellular vesicle-derived miR-425-5p (miR-425) activates astrocytes in the brain to promote breast cancer brain metastasis via the novel miR-425-ZNF24-CCL8 signaling axis

**DOI:** 10.1101/2025.06.05.658130

**Authors:** Grace L. Wong, Munazza S. Khan, Sara Manore, Shivani Bindal, Ravi Singh, Hui-Wen Lo

**Affiliations:** Vivian L. Smith Department of Neurosurgery, McGovern Medical School, The University of Texas Health Science Center at Houston, Houston, Texas, United States; Graduate School of Biomedical Sciences, Wake Forest School of Medicine, Winston-Salem, North Carolina, United States; The University of Texas MD Anderson Cancer Center UTHealth Houston Graduate School of Biomedical Sciences, The University of Texas Health Science Center at Houston, Houston, TX, United States; Department of Cancer Biology, Wake Forest School of Medicine, Winston-Salem, North Carolina, United States; Department of Cell Biology and Genetics, College of Medicine, Texas A&M University Health Science Center at Houston, Bryan, TX, United States

**Keywords:** Breast cancer brain metastasis, microRNAs, extracellular vesicles, brain microenvironment, miR-425

## Abstract

Mechanisms underlying breast cancer brain metastasis (BCBM) are still not well understood. Here, we identified that BCBM patient serum contained extracellular vesicles (EVs) with high levels of microRNAs (miRNAs)-107 and -425. Levels of miR-107 and miR-425 were elevated in brain metastases, and the elevation was associated with poor patient prognoses. Ectopic expression of miR-107 and miR-425 promoted mammospheres; inhibition of miR-425, but not miR-107, suppressed breast cancer mammosphere formation. EVs from miR-425-overexpressing breast cancer cells strongly activated astrocytes whereas their inhibitors abrogated the effect. Conditioned media from miR-425-activated astrocytes promoted mammospheres. Within astrocytes, miR-425 suppressed expression of transcription factor ZNF24, which downregulated CCL8 cytokine expression/secretion, leading to subsequent activation of astrocytes. We further determined the role of miR-425 in brain metastasis formation and observed that miR-425-overexpressing breast cancer cells exhibited significantly more aggressive growth in mouse brains compared to control cells. Immunohistochemistry and immunofluorescence analysis of mouse brain metastases revealed that miR-425 tumors exhibited significantly increased activation, intratumoral accumulation, and proliferation of astrocytes, and a decrease in ZNF24 expression compared to control tumors. Together, our findings demonstrate that breast cancer EV-derived miR-425 promotes BCBM via activating astrocytes in the brain microenvironment through the novel EV-miR-425-ZNF24-CCL8 signaling axis.

## INTRODUCTION

Breast cancer is the most frequently diagnosed cancer and second most common source of brain metastasis in women [1, 2]. It is estimated that 10-40% of patients with solid tumors will develop brain metastases; brain metastases occur almost 10 times more than primary brain tumors [2–5]. Currently, there are no effective treatments to prevent the development of brain metastases from primary breast tumors [6]. Mechanisms that drive breast cancer brain metastasis (BCBM) remain unclear, contributing to dismal survival rates of 6-18 months after diagnosis [7]. Upon diagnosis, HER2-enriched breast cancer and triple-negative breast cancer (TNBC) subtypes display a significantly increased brain metastasis incidence at 8-11%. Furthermore, an estimated 50% of patients develop BCBM over their clinical course, underscoring the importance of identifying the mechanisms that drive breast cancer metastasis to the brain [8–11].

Primary tumors, such as those from the breast, can secrete extracellular vesicles (EVs) to prime pre-metastatic niches in an organotropic-manner [12–14]. EVs transport a variety of bioactive molecules, such as DNA, cytokines, and microRNAs (miRNAs) [15]. MiRNAs are endogenous, small non-coding RNAs that regulate many physiological processes by targeting mRNA transcripts and altering gene expression [15, 16]. Our lab has previously reported that breast cancer-derived EVs containing oncogenic miR-1290 can activate astrocytes to promote tumor growth in the brain, which is also reported for other EV-miRNAs [17–19]. We investigated additional EV-miRNAs and identified miR-425 as a potential regulator of BCBM.

miR-425 expression is increased in breast cancer cells and tissues and is associated with distant metastasis and poor prognoses for breast cancer patients of multiple subtypes [20, 21]. miR-425 plays a major role in the PI3K/AKT signaling pathway by suppressing tumor suppressor PTEN in breast [20], lung [22], and colorectal cancers [23]. Furthermore, overexpression of miR-425 promotes breast cancer tumor growth and metastasis [24]. Whether miR-425 promotes breast cancer metastasis to the brain has not been reported.

In the present study, we examine whether breast cancer-derived EV-miR-425 promotes BCBM by activating astrocytes, the predominant glial cell type in the brain. We found that miR-425 is elevated in BCBM patient serum and brain metastases, with miR-425 activity correlating with worse brain-MFS. Breast-cancer derived EV-miR-425 activates astrocytes and, in turn, miR-425-activated astrocytes promote mammospheres of breast cancer cells—an indicator of cancer stem cell-like properties.

Moreover, we identified Zinc Finger Protein 24 (ZNF24), a miR-425 target, to directly suppress C-C Motif Chemokine Ligand 8 (CCL8), which activates astrocytes and promotes mammosphere formation. Furthermore, HER2-enriched breast cancer cells overexpressing miR-425 enhance the growth of brain metastases, but not bone metastases, *in vivo*. In this study, we discovered a novel role for EV-miR-425 in activating astrocytes in the brain microenvironment, and report that the novel miR-425-ZNF24-CCL8 signaling axis promotes the progression of brain metastases.

## RESULTS

### Breast cancer EV-derived miR-107 and miR-425 are enriched in BCBM patient serum and brain metastases, and are associated with poor breast cancer patient prognoses

We previously observed that breast cancer EV-derived miR-1290, upregulated by BCBM-promoting transcription factor truncated glioma-associated oncogene homolog 1 (tGLI1), promotes the growth of breast cancer cells in the brain by activating astrocytes [17]. Given that the development of BCBM is very complex, additional EV-derived miRNAs could be regulators of BCBM. To identify further breast cancer-specific EV-miRNAs, we analyzed three publicly available Gene Expression Omnibus (GEO) breast cancer patient datasets (GSE68373, GSE37407, and GSE134108; **Fig. 1A**). We found three miRNAs: miR-107, miR-425, and miR-1290 that were significantly upregulated in the serum of breast cancer patients with metastases compared to those without (GSE68373), enriched in brain metastases compared to matched primary breast tumors (GSE37407), and increased in the serum of patients with BCBM compared to those without metastases (GSE134108; **Fig. 1B**). Since we previously published that miR-1290 promotes BCBM, we decided to focus on the potential roles of miR-107 and miR-425 in BCBM. We found that miR-107 and miR-425 are significantly increased in serum from patients with breast cancer compared to non-cancer controls (GSE73002; **Fig. 1C**). Additionally, miR-107 and miR-425 are upregulated in primary breast tumors compared to matched, normal adjacent tissues (GSE40525; **Fig. 1D**). To determine whether these miRNAs could predict patient prognoses, we analyzed miRNA expression and gene signatures [25] in GEO patient datasets. We found that miR-107 and miR-425 expression correlates with worse distant relapse-free survival (GSE22220; **Fig. 1E**). In addition, the predicted activity of both miRNAs correlates with shortened metastasis-free survival (MFS) and brain-MFS of breast cancer patients (GSE12276/2034/2603/5327/14020; **Fig. 1F-G**). We also found that genes associated with brain metastatic behavior (“Breast-to-Brain” GS) [26] are significantly positively enriched with the miR-425 gene signature; the “Breast-to-Brain” GS is enriched with the miR-107 gene signature, but did not reach statistical significance (**Fig. 1H**). Next, we isolated EVs, followed by miRNAs from the serum of breast cancer patients treated at the Wake Forest Baptist Comprehensive Cancer Center (WFBCCC), which revealed that both miR-107 and miR-425 are upregulated in serum samples from patients with brain metastases (N = 7) versus those with no metastases (N = 11; **Fig. 1I**). Taken together, these data demonstrate that miR-107 and miR-425 may play important roles in BCBM.

**Figure 1:**
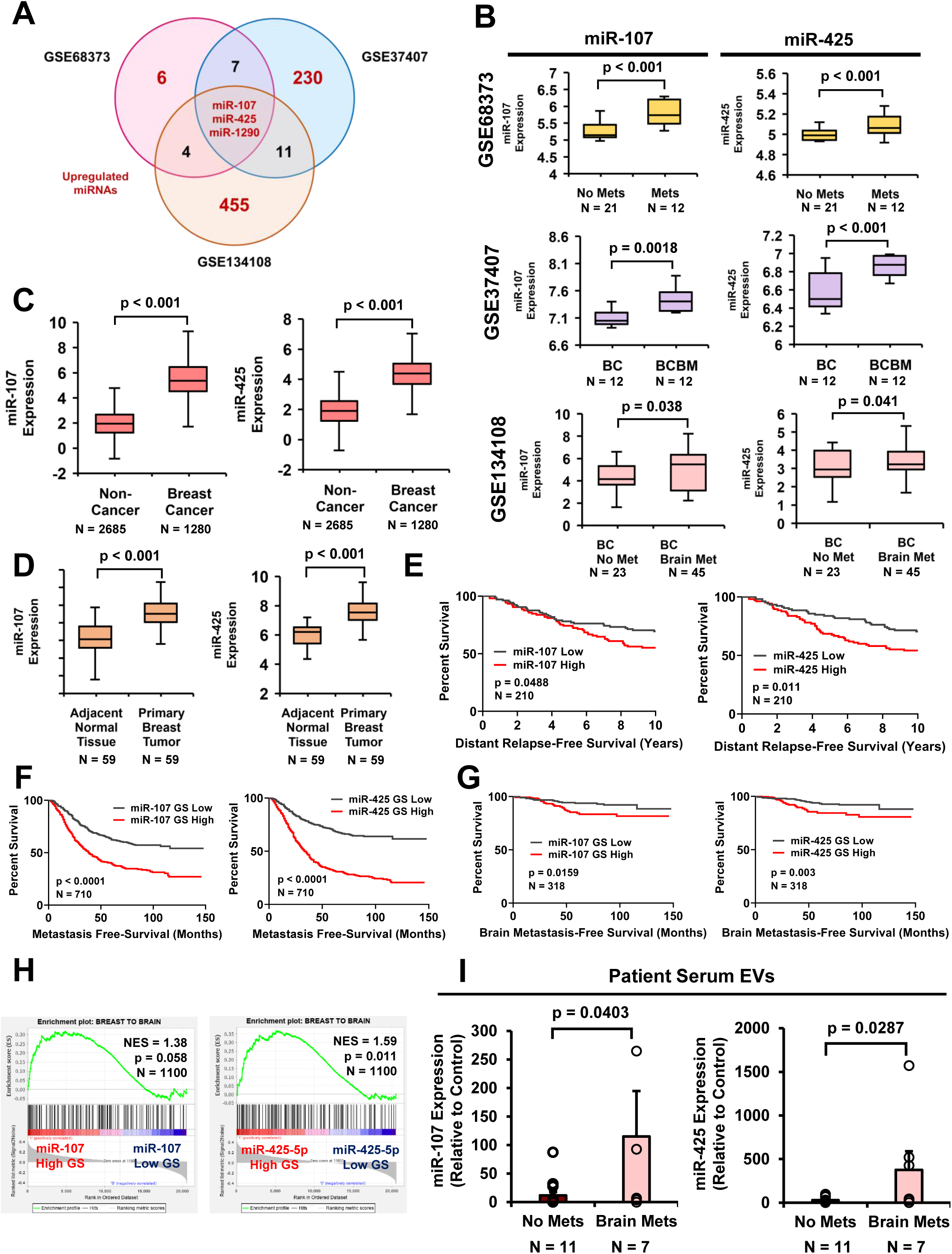
Breast cancer EV-derived miR-107 and miR-425 are enriched in BCBM patient serum and brain metastases, and are associated with poor breast cancer patient prognoses. **A-B)** Identification of three miRNAs that were upregulated in 1) serum of patients with metastases compared to no metastases (GSE68373), 2) brain metastases compared to matched primary tumors (GSE37407), and 3) serum of patients with only brain metastases compared to no metastases (GSE134108). **C)** miR-107 and miR-425 are enriched in serum samples of breast cancer patients compared to non-cancer controls (GSE73002). **D)** miR-107 and miR-425 are enriched in primary breast tumors compared to matched, normal adjacent tissues (GSE40525). **E)** Expression of miR-107 and miR-425 correlates with worse distant relapse-free survival (GSE22220). **F-G)** The miR-107 and miR-425 gene signatures (GS) correlate with shortened metastasis-free survival (MFS) and brain MFS in breast cancer patients (GSE12276/2034/2603/5327/14020). **H)** miR-107 and miR-425 GS are enriched with genes upregulated in breast cancer brain metastasis [26] in the Breast Cancer TCGA dataset using GSEA analysis. **I)** miR-107 and miR-425 expression is upregulated in serum from patients with only brain metastases compared to serum from patients with no metastases (WFBCCC patient serum). Fold change was calculated in Panels B-D, and I. Student’s *t*-test was used in Panels B-D, and I. Log-Rank (Mantle-Cox) was used in Panels E-G. N = 3 experimental replicates unless otherwise indicated.

### miR-107 and miR-425 are upregulated in EVs from brain-tropic cells and miR-425 promotes mammosphere formation of breast cancer cells

Given that miR-107 and miR-425 are upregulated in BCBM patient serum and brain metastases (**Fig. 1**), we wanted to examine expression in brain-tropic breast cancer cell lines. We analyzed EVs from the conditioned medium (CM) of parental HER2-enriched breast cancer and TNBC cell lines, SKBR3 and CN34, and their brain-metastatic sublines, SKBRM and CN34-BRM, respectively. Both miR-107 and miR-425 are enriched in EVs from brain-tropic cells (SKBRM and CN34-BRM) compared to the parental cells (SKBR3 and CN34; **Fig. 2A-B**). EVs were isolated using the ExoQuick-TC ULTRA Kit (System Biosciences) and validated via Nanoparticle Tracking Analysis (NTA; **Supplementary Fig. 1A**). Consistent with EV-miRNA findings, intracellular miR-425 levels are increased in brain-tropic cells compared to parental breast cancer cells; intracellular miR-107 levels are increased in CN34-BRM cells compared to CN34 cells (**Fig. 2C-D**). To further investigate the roles of miR-107 and miR-425 in breast cancer cells, we utilized miR-107 and miR-425 mimics and inhibitors (ThermoFisher Scientific) and validated overexpression with miR-RT-qPCR in SKBR3 cells (**Fig. 2E**). Next, we wanted to examine whether these two miRNAs could promote malignant phenotypes that are characteristic of breast cancer metastasis. Mammosphere assays enrich the breast cancer stem cell (BCSC) subpopulation responsible for breast cancer metastasis [27]. Interestingly, transfection with the miR-425 mimic, not the miR-107 mimic, promoted mammosphere formation of SKBR3 cells (**Fig. 2F**). Moreover, the inhibition of miR-425, not miR-107, suppressed mammosphere-forming ability of SKBRM cells (**Fig. 2G**). Inhibition of miR-107 and miR-425 was validated by measuring mRNA levels of previously published target genes where we expect that the successful inhibition of each miRNA will increase target gene expression [22, 28–34]. We found that only one of the miR-107 target genes significantly increased with miR-107 inhibition, while four out of seven miR-425 target genes significantly increased (**Supplementary Fig. 1C-D**). Given that most of the miR-107 target genes did not increase with miR-107 inhibition, knockdown of miR-107 alone may not be sufficient to suppress its target genes. This observation can be explained by the fact that miR-107 is a member of the miR-15/103/107 gene family, meaning these miRNAs share similar seed regions and can downregulate the same genes [35].

**Figure 2:**
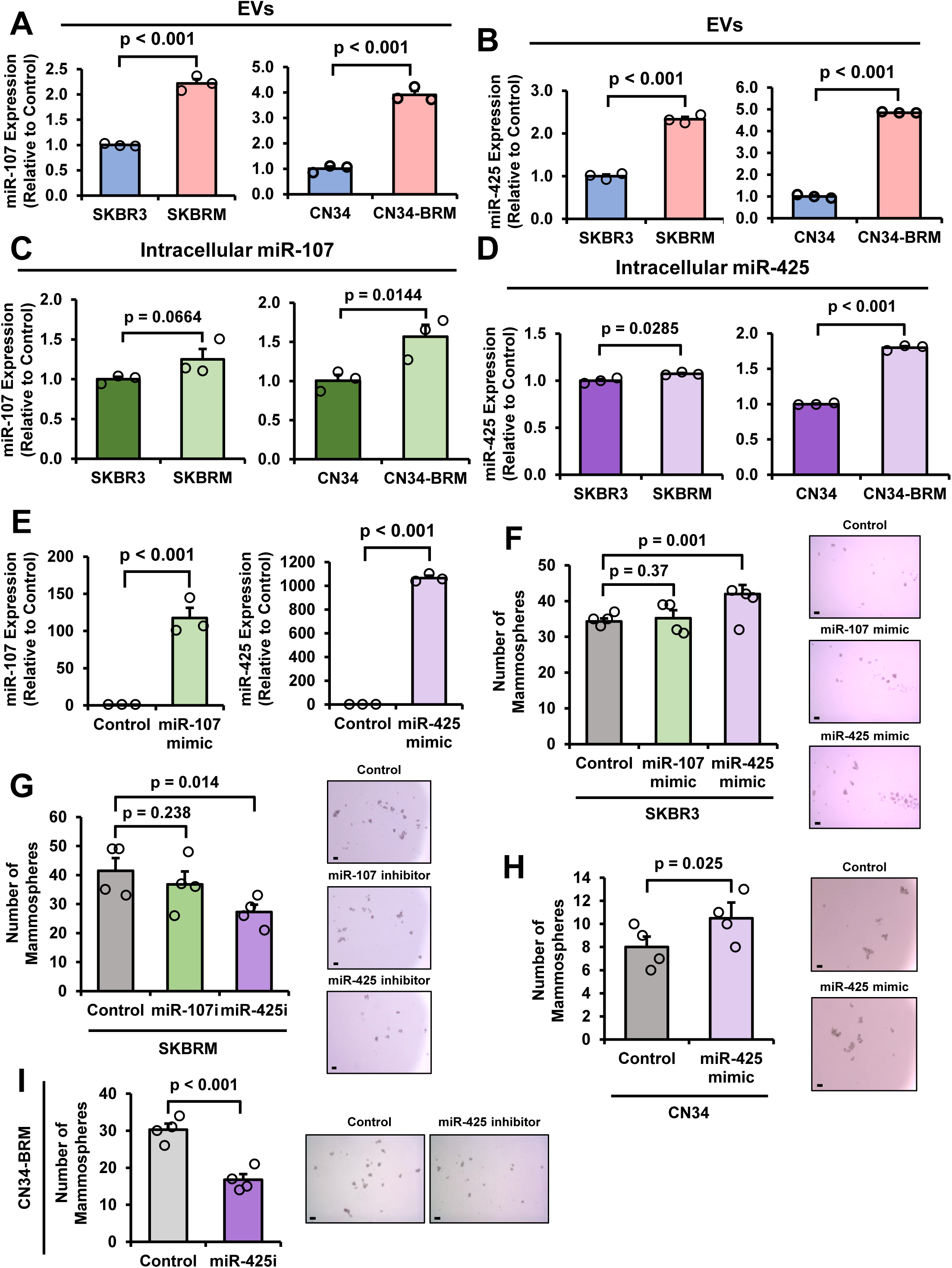
miR-107 and miR-425 are upregulated in EVs from brain-tropic cells and miR-425 promotes mammosphere formation of breast cancer cells. **A-B)** miR-107 and miR-425 are enriched in EVs from brain-tropic cells (SKBRM and CN34-BRM) compared to the parental cells (SKBR3 and CN34). **C-D)** Intracellular miR-107 and miR-425 expression are increased in brain-tropic cells compared to parental cells. **E)** Overexpression of miR-107 and miR-425 with miRNA mimics in SKBR3 cells validated via miR-RT-qPCR. **F)** Ectopic expression of miR-425, not miR-107, promotes mammosphere formation of SKBR3 cells. Scale bar indicates 100 µm. **G)** Inhibition of miR-425, not miR-107, suppresses the mammosphere-forming ability of SKBRM cells. Scale bar indicates 100 µm. **H)** Ectopic expression of miR-425 promotes mammosphere formation of CN34 cells. Scale bar indicates 100 µm. **I)** miR-425 inhibition suppresses mammosphere formation of CN34-BRM cells. Scale bar indicates 100 µm. Fold change was calculated in Panels A-E. Student’s *t*-test was used in Panels A-I. N = 3 experimental replicates unless otherwise indicated.

Additionally, miR-107 has been reported to play a tumor suppressive role in breast cancer [36]. These findings directed our focus to miR-425 and its role in BCBM. Consistent with our findings in SKBR3 and SKBRM cells, overexpression of miR-425 promoted mammosphere formation of CN34 cells, while miR-425 inhibition suppressed mammosphere-forming ability of CN34-BRM cells (**Fig. 2H-I**). Validation of miR-425 overexpression in CN34 cells was measured by miR-RT-qPCR (**Supplementary Figure 1B**). Collectively, these findings indicate that miR-425 contributes to breast cancer mammospheres and, potentially, BCBM.

### miR-425 activates astrocytes and miR-425-activated astrocytes promote mammospheres

We found EVs from BCBM patients are enriched with miR-425 (**Fig. 1I**), so next we wanted to investigate the mechanism by which these EV-miRNAs could alter brain microenvironmental cells to promote BCBM. Often referred to as the “pre-metastatic niche”, cancer cells prime brain microenvironmental cells to promote tumor growth in the brain by utilizing a variety of methods including secreting EVs to activate brain microenvironmental cells, such as astrocytes. Astrocytes, specialized glial cells, play critical roles in supporting neurons, immune signaling, providing structural support and organization, regulating central nervous system (CNS) blood flow, and are one of the three main components that make up the blood-brain barrier (BBB) [37, 38]. Astrocytes are highly abundant in the brain and reactive astrocytes (also referred to as activated astrocytes) promote brain metastasis by stimulating the release of tumor-supportive cytokines and molecules, inhibiting tumor suppressive proteins, promoting tumor-astrocyte gap junctions, and protecting tumor cells from toxic chemotherapy [6, 13, 17, 37, 39–41]. To determine whether miR-425 activates astrocytes, we transfected human astrocytes with the miR-425 mimic and performed glial fibrillary acidic protein (GFAP) immunofluorescence (IF), which is indicative of astrocyte activation [13]. Ectopic expression of miR-425 in human astrocytes significantly increased GFAP protein and mRNA expression (**Fig. 3A-B**).

**Figure 3:**
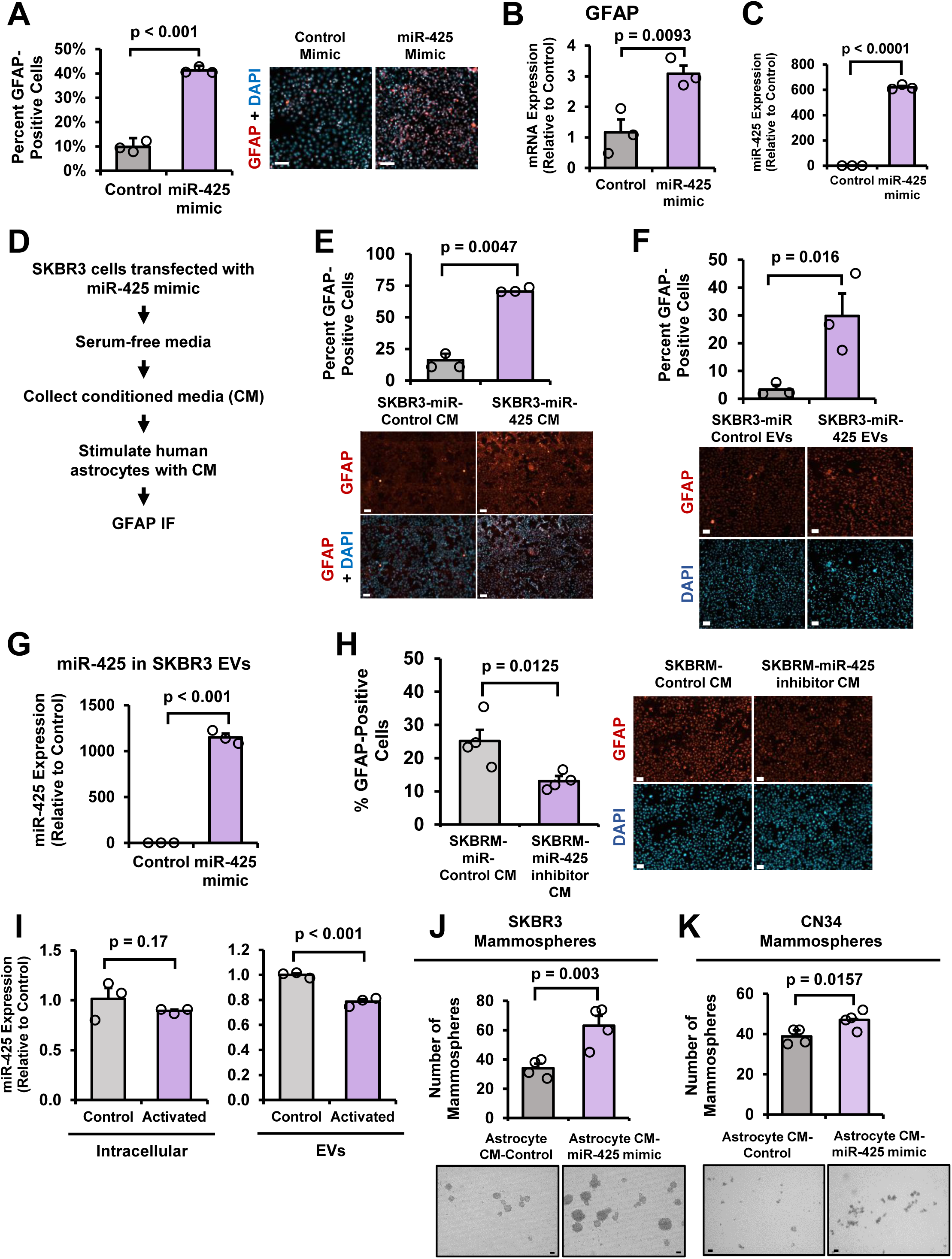
miR-425 activates astrocytes and miR-425-activated astrocytes promote mammospheres. **A)** Overexpression of miR-425 in human astrocytes significantly increases GFAP expression as indicated by GFAP immunofluorescence (IF). Representative 20x images. Scale bar indicates 100 µm. **B)** Transfection of miR-425 in human astrocytes significantly increases GFAP mRNA expression. **C)** Validation of miR-425 overexpression in human astrocytes via miR-RT-qPCR. **D-E)** CM from SKBR3 cells with miR-425 mimic increases GFAP expression as measured by GFAP IF. Representative 20x images. Scale bar indicates 100 µm. **F)** EVs from SKBR3 cells overexpressing miR-425 significantly increase GFAP expression in human astrocytes as measured by GFAP IF. Representative 20x images. Scale bar indicates 100 µm. **G)** Validation of miR-425 overexpression in EVs from SKBR3 cells transfected with the miR-425 mimic as measured by miR-RT-qPCR. **H)** miR-425 inhibition in SKBRM cells suppresses GFAP expression in human astrocytes. Representative 20x images. Scale bar indicates 100 µm. **I)** Intracellular and EV-miR-425 expression in astrocytes activated by a cocktail of IL-1α, TNFα, and C1q. **J-K)** CM from astrocytes overexpressing miR-425 promotes mammosphere formation of SKBR3 and CN34 cells. Scale bar indicates 100 µm. Fold change was calculated in Panels B-C, G, and I. Student’s *t*-test was used in Panels A-C, E-K. N = 3 experimental replicates unless otherwise indicated.

Overexpression of miR-425 in the human astrocytes was validated via miR-RT-qPCR (**Fig. 3C**). Given that brain-tropic breast cancer-derived EVs contain elevated miR-425 levels, we transfected SKBR3 cells with the miR-425 mimic, collected the CM, and stimulated the human astrocytes with CM. We found that overexpression of miR-425 in SKBR3 cells significantly activated astrocytes (**Fig. 3D-E**). To determine whether the EVs from the breast cancer cells are responsible for the astrocyte activation, we isolated EVs from SKBR3 cells overexpressing miR-425, and stimulated astrocytes with the EVs.

Analysis of GFAP IF reveals that astrocytes stimulated with EVs from SKBR3 cells overexpressing miR-425 are significantly more activated than EVs from the control cells (**Fig. 3F**). Overexpression of miR-425 in the EVs is validated via miR-RT-qPCR (**Fig. 3G**). To complement our overexpression assays, SKBRM cells were transfected with the miR-425 inhibitor and the CM was used to stimulate astrocytes. Inhibition of miR-425 in SKBRM cells suppresses GFAP expression in astrocytes (**Fig. 3H**). Since overexpression of miR-425 and CM from SKBR3 cells overexpressing miR-425 activates astrocytes, we wanted to determine whether activated astrocytes also increase expression of miR-425. We stimulated astrocytes with a previously published cocktail containing recombinant human IL-1α, TNFα, and C1q complement protein [42] and analyzed miR-425 expression using miR-RT-qPCR. Interestingly, we found that both intracellular and EV-miR-425 levels were decreased in astrocytes activated by the recombinant human IL-1α, TNFα, and C1q complement protein cocktail, suggesting that miR-425 expression is derived from the breast cancer cells, not the astrocytes (**Fig. 3I**). To examine whether astrocytes activated by miR-425 could promote BCSCs, we collected CM from astrocytes transfected with the miR-425 mimic and stimulated breast cancer cells in mammosphere assays. Astrocytes activated by miR-425 significantly increase mammosphere formation of SKBR3 and CN34 cells (**Fig. 3J-K**). Taken together, these results demonstrate that breast cancer cells overexpressing miR-425 secrete EVs with high levels of miR-425 that can activate astrocytes, and those miR-425-activated astrocytes promote mammospheres of TNBC and HER2-enriched breast cancer cells.

### miR-425-activated astrocytes secrete CCL8 and SCF to further activate astrocytes and promote mammospheres

Next, we wanted to examine the mechanism by which CM from miR-425-activated astrocytes promotes mammosphere formation of SKBR3 and CN34 cells (**Fig. 3J-K**). We reasoned that miR-425-activated astrocytes may secrete specific cytokines to promote breast cancer cells. To identify these cytokines, we collected CM from astrocytes transfected with the control or miR-425 mimic and performed a cytokine array comprising over 80 cytokines (**Fig. 4A**). Multiple cytokines were up-or down-regulated in CM from miR-425-activated astrocytes; we focused on the top 4 upregulated cytokines: CCL8 (also known as MCP-2), Interleukin-4 (IL-4), Stem cell factor (SCF, gene name is KITLG), and Macrophage migration inhibitory factor (MIF). Reactive astrocytes often secrete cytokines to activate nearby astrocytes [17], so next we examined whether these cytokines could activate astrocytes. We found that all four cytokines could independently significantly increase GFAP expression in astrocytes (**Fig. 4B**).

**Figure 4:**
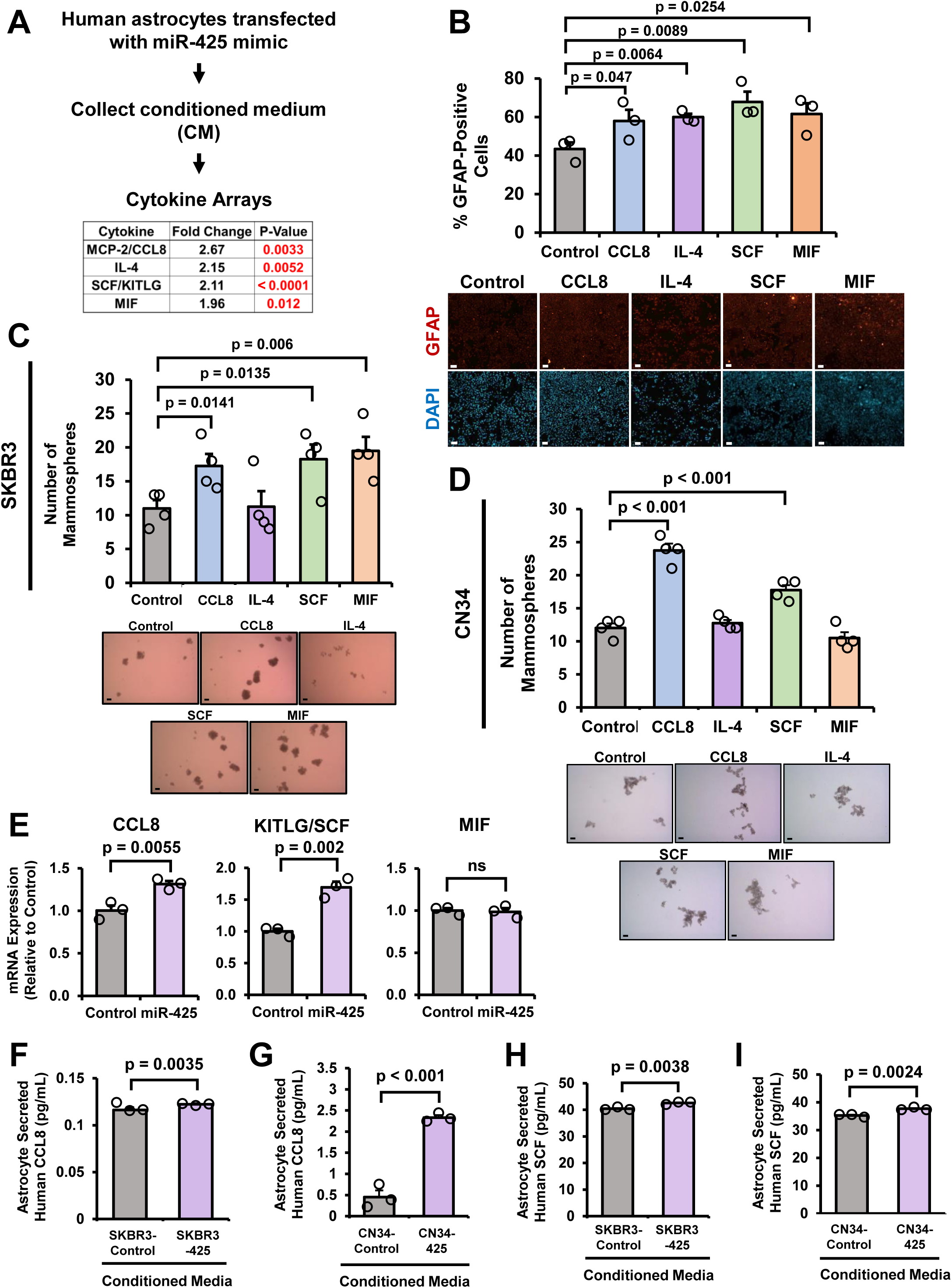
miR-425-activated astrocytes secrete CCL8 and SCF to further activate astrocytes and promote mammospheres. **A)** Top four highly secreted cytokines in CM from astrocytes overexpressing miR-425 as measured by cytokine arrays. **B)** Cytokine stimulation increases GFAP expression in human astrocytes as indicated by GFAP IF. Representative 20x images. Scale bar indicates 100 µm. **C)** CCL8, SCF, and MIF stimulation promotes SKBR3 mammospheres. Scale bar indicates 100 µm. **D)** CCL8 and SCF promote mammosphere formation of CN34 cells. Scale bar indicates 100 µm. **E)** CCL8 and KITLG are upregulated in astrocytes overexpressing miR-425 as measured by RT-qPCR. **F)** Human astrocytes stimulated by CM from SKBR3 cells overexpressing miR-425 secrete significantly higher levels of CCL8 compared to astrocytes stimulated with CM from control SKBR3 cells. Secreted CCL8 expression measured by ELISA. **G)** Astrocytes stimulated by CM from CN34 overexpressing miR-425 secrete significantly increased levels of CCL8 compared to astrocytes stimulated with CM from control CN34 cells. Secreted CCL8 expression measured by ELISA. **H-I)** Astrocytes stimulated by CM from SKBR3 or CN34 cells overexpressing miR-425 secrete high levels of SCF. Secreted SCF expression measured by ELISA. Fold change was calculated in Panels E. Student’s *t*-test was used in Panels B-I. N = 3 experimental replicates unless otherwise indicated.

Since miR-425-activated astrocytes promote breast cancer mammosphere formation, we wanted to determine whether these cytokines promote mammospheres. We stimulated SKBR3 and CN34 cells with recombinant protein and cultured the cells under mammosphere-forming conditions. Our results reveal that CCL8, SCF, and MIF significantly promote mammosphere-forming ability in SKBR3 cells, while only CCL8 and SCF significantly increase mammospheres in CN34 cells (**Fig. 4C-D**). To further narrow down these three cytokines, we compared mRNA expression of CCL8, KITLG (SCF), and MIF in astrocytes overexpressing miR-425. CCL8 and KITLG (SCF) mRNA levels are significantly increased in astrocytes overexpressing miR-425 (**Fig. 4E**). RT-qPCR analysis revealed that CCL8, SCF, and MIF receptor expression remained unchanged in human astrocytes transfected with the miR-425 mimic, suggesting that miR-425 upregulates CCL8 or SCF cytokine expression and not the corresponding receptor (**Supplementary Fig. 2A**). To further validate the cytokine array results, we stimulated astrocytes with CM from SKBR3 cells transfected with the control or miR-425 mimic. CM from SKBR3 cells overexpressing miR-425 significantly increased the ability of astrocytes to secrete CCL8 compared to CM from control cells (**Fig. 4F**). Furthermore, CM from CN34 cells transfected with the miR-425 mimic significantly increased CCL8 secretion by astrocytes (**Fig. 4G**). We also found increased human SCF secretion from astrocytes stimulated by SKBR3 or CN34 cells overexpressing miR-425 (**Fig. 4H-I**). These findings suggest that miR-425-activated astrocytes secrete high levels of CCL8 and SCF that further activate astrocytes and promote mammosphere formation.

### miR-425 suppresses transcription factor ZNF24 in astrocytes

Given that miRNAs regulate gene expression by suppressing mRNA transcripts, we aimed to determine the mechanism by which miR-425-activated astrocytes secrete high levels of CCL8 and SCF. To do this, we needed to identify a transcriptional repressor of CCL8 or SCF that is a target of miR-425. First, we identified transcription factors with predicted binding sites within the CCL8 and SCF promoter regions on GeneCards [43]. Next, we examined miR-425 binding sites within 3’-untranslated regions (UTRs) on TargetScan [44] and found BCOR, CREB1, and ZNF24 filled both criteria: 1) contained predicted binding sites within the CCL8 or SCF promoter regions, and 2) were predicted miR-425 target genes (**Fig. 5A**). To validate these findings, we compared mRNA levels in astrocytes overexpressing miR-425 with the expectation that direct miR-425 targets will have decreased mRNA expression.

**Figure 5:**
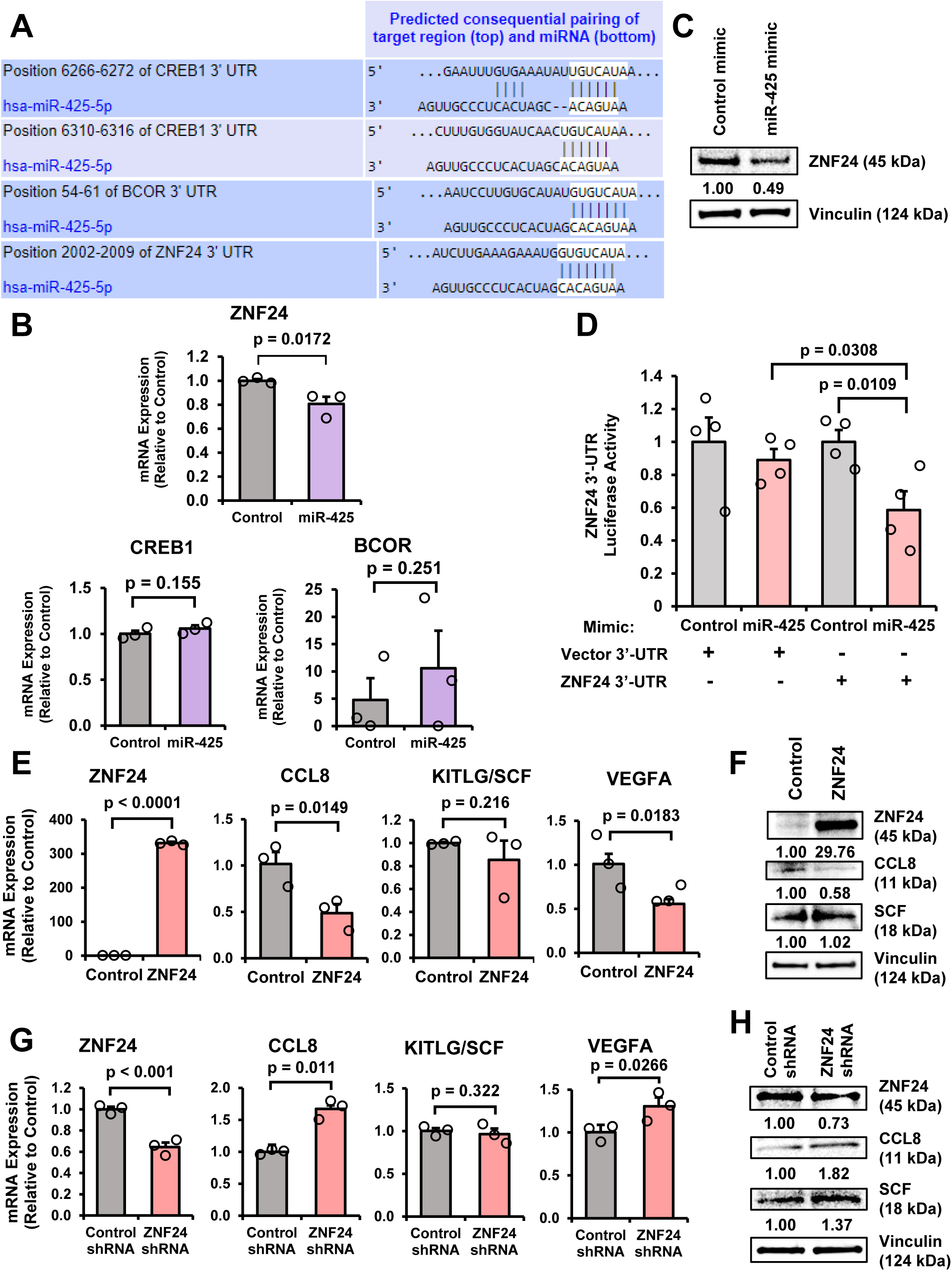
miR-425 suppresses transcription factor ZNF24 in astrocytes. **A)** Predicted miR-425 binding sites within the CREB1, BCOR, and ZNF24 3’-UTRs from TargetScan. **B)** ZNF24 mRNA expression is significantly decreased in astrocytes overexpressing miR-425. CREB1 and BCOR mRNA expression is unchanged as measured by RT-qPCR. **C)** ZNF24 protein expression is decreased in astrocytes transfected with miR-425 mimic as indicated by western blot analysis. **D)** miR-425 suppresses ZNF24 3’-UTR activity as measured by dual luciferase reporter assay. **E)** Ectopic expression of ZNF24 significantly decreases CCL8 and VEGFA mRNA expression, while KITLG is unchanged. mRNA expression measured by RT-qPCR. **F)** Ectopic expression of ZNF24 suppresses CCL8 protein expression, but not SCF as indicated by western blot analysis. **G)** ZNF24 knockdown significantly increases CCL8 and VEGFA mRNA expression, but not KITLG. mRNA expression as indicated by RT-qPCR. **H)** ZNF24 knockdown increases CCL8 and SCF protein expression as measured by western blot analysis. Fold change was calculated in Panels B, D, E, and G. Student’s *t-*test was used in Panels B, D, E, and G. N = 3 experimental replicates unless otherwise indicated.

Interestingly, only Zinc Finger Protein 24 (ZNF24) is significantly suppressed in astrocytes overexpressing miR-425 (**Fig. 5B**). Moreover, western blot analysis revealed that ZNF24 protein expression decreased with miR-425 overexpression (**Fig. 5C**). Furthermore, using a luciferase reporter controlled by the ZNF24 3’-UTR, we found that transfection with the miR-425 mimic significantly reduced activity of the ZNF24 3’-UTR reporter compared to the control mimic and vector plasmid, indicating miR-425 binds to the ZNF24 3’-UTR (**Fig. 5D**). To determine whether ZNF24 suppressed CCL8 and SCF (KITLG) expression, we utilized a plasmid designed to overexpress ZNF24 (VectorBuilder) and examined CCL8, SCF (KITLG), and VEGFA expression. VEGFA is directly suppressed by ZNF24 [45] and is a positive control. As predicted, ectopic expression of ZNF24 increased ZNF24 and decreased CCL8 mRNA and protein expression (**Fig. 5E-F**). VEGFA mRNA expression is suppressed, while KITLG mRNA is unchanged. To complement these overexpression studies, we transfected astrocytes with ZNF24 short-hairpin RNA (shRNA) and found that ZNF24 mRNA and protein expression decreases, while CCL8 mRNA and protein expression increase (**Fig. 5G-H**). RT-qPCR analysis revealed an increase in VEGFA mRNA levels and SCF (KITLG) mRNA levels remained unchanged. SCF protein expression increases with ZNF24 shRNA, suggesting that ZNF24 may not directly suppress SCF expression. These findings demonstrate that the potential miR-425-ZNF24-CCL8 signaling axis may play an important role in reactive astrocytes.

### ZNF24 suppresses astrocyte activation and directly decreases CCL8 expression

Given that miR-425 is a direct repressor of ZNF24 (**Fig. 5D**), we aimed to determine whether ZNF24 could regulate astrocyte activation. To do this, we first activated astrocytes with a previously published cocktail containing recombinant human IL-1α, TNFα, and C1q complement protein [42] and validated the significant increase in GFAP protein and mRNA expression (**Fig. 6A-B**). Next, we activated astrocytes, then transfected activated astrocytes with control or ZNF24 plasmids and measured GFAP protein and mRNA expression. Our results demonstrate that ectopic expression of ZNF24, validated by ZNF24 mRNA levels, suppresses GFAP protein and mRNA expression (**Fig. 6C-D**). To determine whether ZNF24 directly binds to the CCL8 promoter, we performed ChIP-qPCR and observed that ZNF24 directly binds to two regions within the human CCL8 promoter (**Fig. 6E**). Furthermore, using a luciferase reporter under the control of the CCL8 promoter, we found that ZNF24 suppresses CCL8 promoter transactivation compared to vector (**Fig. 6F**). To determine whether miR-425 activates this pathway in breast cancer cells, we transfected breast cancer cells with the miR-425 mimic and performed RT-qPCR for ZNF24 and CCL8 expression. Interestingly, ZNF24 and CCL8 mRNA expression is not significantly different in breast cancer cells overexpressing miR-425 suggesting this miR-425-ZNF24-CCL8 signaling axis is specific to astrocytes (**Fig. 6G-H**). Taken together, our results demonstrate, for the first time, that ZNF24 suppresses astrocyte activation and that ZNF24 directly binds and represses CCL8 expression in astrocytes.

**Figure 6:**
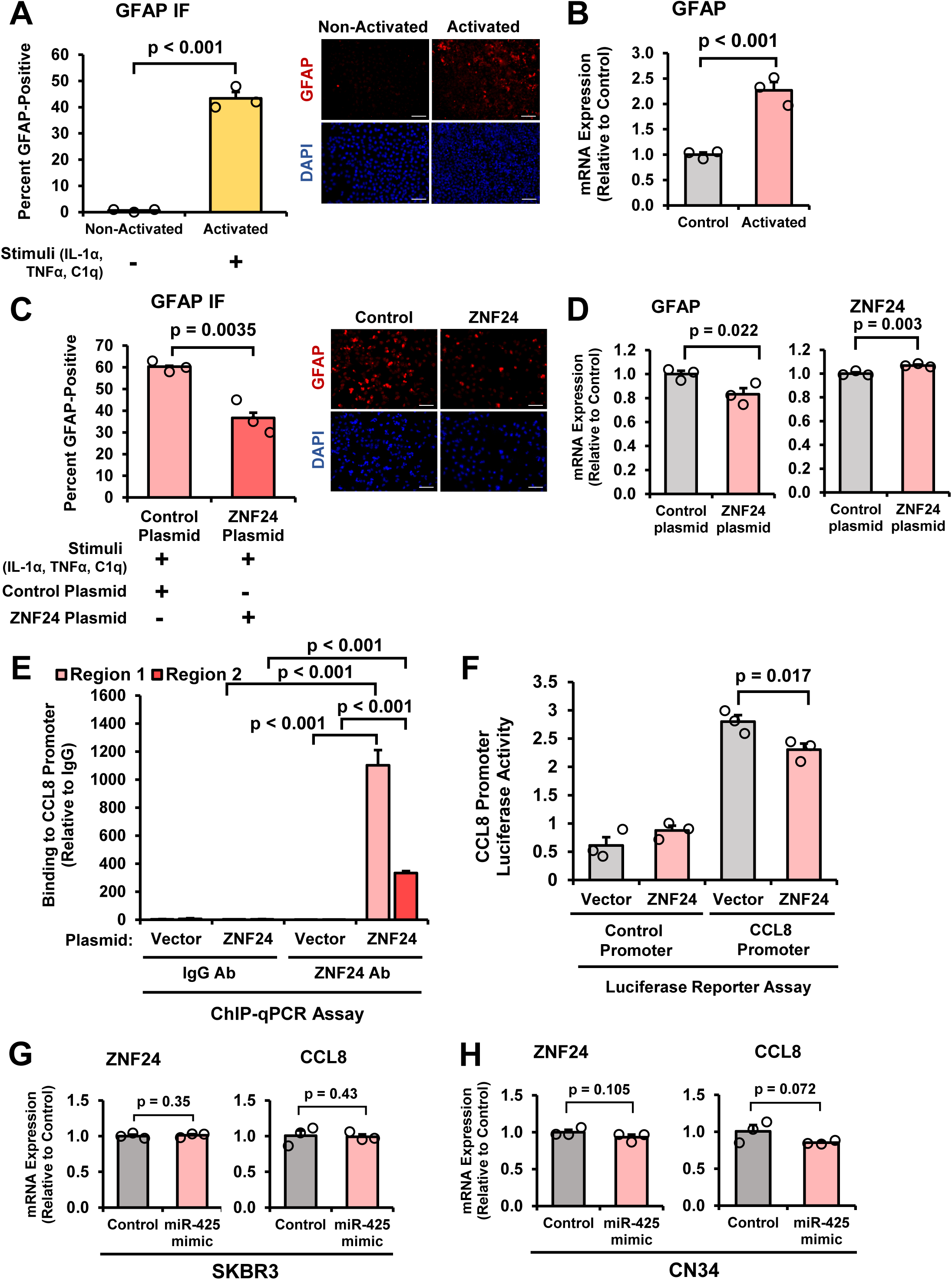
ZNF24 suppresses astrocyte activation and decreases CCL8 expression. **A)** Human astrocytes activated by a cocktail of IL-1α, TNFα, and C1q. Representative 20x images. Scale bar indicates 100 µm. **B)** Validation of GFAP mRNA expression in activated astrocytes as indicated by RT-qPCR. **C)** Activated astrocytes transfected with ZNF24 plasmid have significantly reduced GFAP expression as indicated by GFAP IF. Representative 20x images. Scale bar indicates 100 µm. **D)** Validation of GFAP reduction and ZNF24 overexpression with ectopic expression of ZNF24. mRNA expression measured by RT-qPCR. **E)** ZNF24 binding to the CCL8 promoter in two regions (−69 and -205 bases upstream of the TSS) measured by ChIP-qPCR. **F)** Ectopic expression of ZNF24 suppresses CCL8 promoter activity as indicated by dual luciferase reporter assays. **G-H)** ZNF24 and CCL8 mRNA expression remains unchanged in breast cancer cells transfected with control or miR-425 mimic. mRNA expression measured by RT-qPCR. Fold change was calculated in Panels B, D-H. Student’s *t*-test was used in Panels A-H. N = 3 experimental replicates unless otherwise indicated.

### Overexpression of miR-425 promotes the growth of breast cancer cells in mouse brains

Since we observed breast cancer-derived EV-miR-425 activate astrocytes and miR-425-activated astrocytes further activate astrocytes and promote mammospheres, we investigated whether miR-425 could promote BCBM *in vivo*. We lentivirally-infected SKBR3 cells to stably express luciferase and miR-425. Overexpression of miR-425 in SKBR3-Luc-miR-425 cells compared to SKBR3-Luc-miR-Control cells is validated in both cells and EVs (**Fig. 7A**). Furthermore, CM from SKBR3-Luc-miR-425 cells significantly activated astrocytes compared to control (**Fig. 7B**). Whether miR-425 promotes BCBM *in vivo* has not been studied. We intracardially inoculated 5-6 week-old, athymic female mice with SKBR3-Luc-miR-Control and SKBR3-Luc-miR-425 cells and monitored metastases via IVIS imaging twice weekly (**Fig. 7C**). We observed that mice intracardially injected with SKBR3-Luc-miR-425 cells had significantly larger brain metastases compared to those intracardially injected with SKBR3-Luc-miR-Control cells (**Fig. 7D, F**). Interestingly, we found that SKBR3-Luc-miR-Control mice exhibited significantly larger bone metastases compared to the SKBR3-Luc-miR-425 group, supporting the brain-tropism of miR-425 (**Fig. 7E-F**). Furthermore, *ex vivo* IVIS imaging of the brain and bone revealed that the SKBR3-Luc-miR-425 group had significantly larger brain metastases and significantly smaller bone metastases compared to the control group (**Fig. 7G-I**).

**Figure 7:**
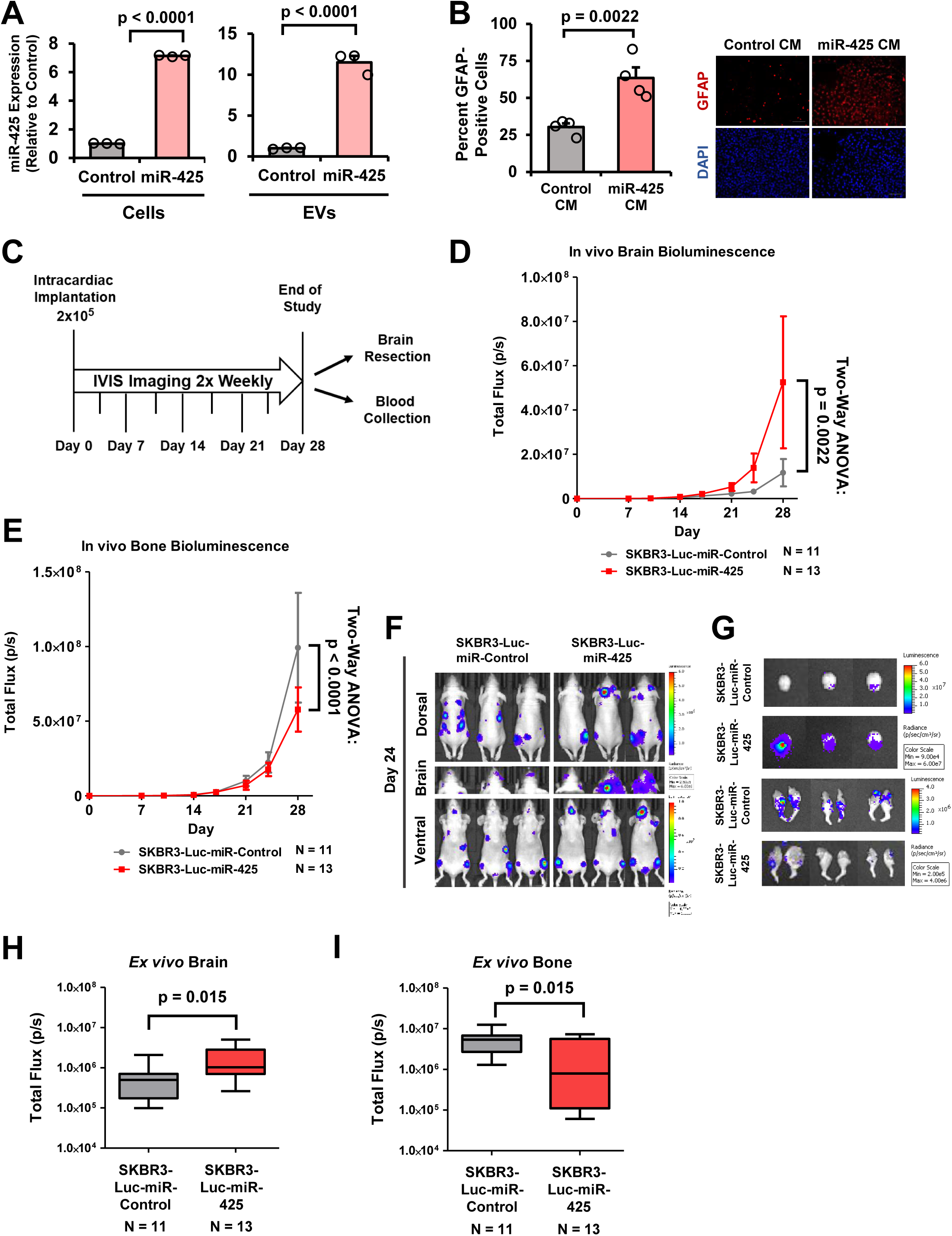
Overexpression of miR-425 promotes the growth of breast cancer cells in mouse brains. **A)** Generation of SKBR3-Luc-miR-Control and -425 stable cell lines. Overexpression of miR-425 in SKBR3-Luc-miR-425 in cells and EVs validated by RT-qPCR. **B)** SKBR3-Luc-miR-425 cells increase GFAP expression in human astrocytes as indicated by GFAP IF. Representative 20x images. Scale bar indicates 100 µm. **C)** Experimental design of intracardiac animal study with SKBR3-Luc-miR-Control and SKBR3-Luc-miR-425 cells in 5–6-week-old female, athymic mice. **D)** Bioluminescence measuring tumor growth in the brain throughout the study. Two-way ANOVA used to compute p-value. **E)** Bioluminescence measuring tumor growth in the bone throughout the study. Two-way ANOVA used to compute p-value. **F)** Representative IVIS images from Day 24. **G)** Representative *ex vivo* IVIS images of brain and bone. **H-I)** Bioluminescence *ex vivo* IVIS of brain and bone. Fold change was calculated in Panel A. Student’s *t*-test was used in Panels A-B, and H-I.

Next, we wanted to examine the miRNA expression in circulating EVs from the intracardiac study. We isolated EVs from mouse serum, then miRNAs, and performed RT-qPCR. We found significantly higher levels of EV-miR-425 in serum from mice in the SKBR3-Luc-miR-425 group compared to the control group (**Fig. 8A**). Since we observed CM from breast cancer cells overexpressing miR-425 to stimulate higher levels of CCL8 secretion in astrocytes, we examined CCL8 secretion in mouse serum. We found serum from SKBR3-Luc-miR-425 mice had significantly increased CCL8 secretion compared to the control group serum (**Fig. 8B**). We found no significant difference in SCF levels in the mouse serum (**Fig. 8C**). Next, we analyzed tissue sections of the mouse brains containing brain metastases; brain metastases were identified based on GFP expression and confirmed with hematoxylin & eosin (H&E) staining. Immunohistochemistry (IHC) analysis reveals the SKBR3-Luc-miR-425 brain metastases had increased tumor-adjacent activated astrocytes (GFAP), increased intratumoral astrocytes (GFAP), and increased breast cancer cell proliferation (Ki-67; **Fig. 8D-F, H**). Furthermore, we found that astrocytes within and surrounding the brain metastases from the SKBR3-Luc-miR-425 group had significantly decreased levels of ZNF24 expression (**Fig. 8G-H**). To further confirm that the ZNF24 expression is specific to the astrocytes, we performed co-staining IF for ZNF24 and GFAP in mouse brain sections containing the brain metastases. Since GFAP expression (red) is used to detect astrocytes, we quantified the percentage of astrocytes with positive ZNF24 expression (green) and found significantly decreased ZNF24 expression in the GFAP-positive astrocytes from the miR-425 group compared to the control (**Fig. 8I**). In a previous report, multiple PTEN-targeting EV-miRNAs are secreted by reactive astrocytes to suppress PTEN expression in only brain metastases [46]. Given that miR-425 is reported to directly suppress PTEN [20], we examined PTEN protein expression in brain metastases from the intracardiac animal model. We found no significant change in PTEN expression in mouse brain metastases indicating that miR-425 is not directly suppressing PTEN in this context (**Supplementary Fig. 3A-B**). In summary, we report that brain-metastatic breast cancer cells secrete increased levels of EV-miR-425, which activates astrocytes through the novel miR-425-ZNF24-CCL8 signaling pathway to promote the growth of breast cancer cells in the brain (**Fig. 8J**).

**Figure 8:**
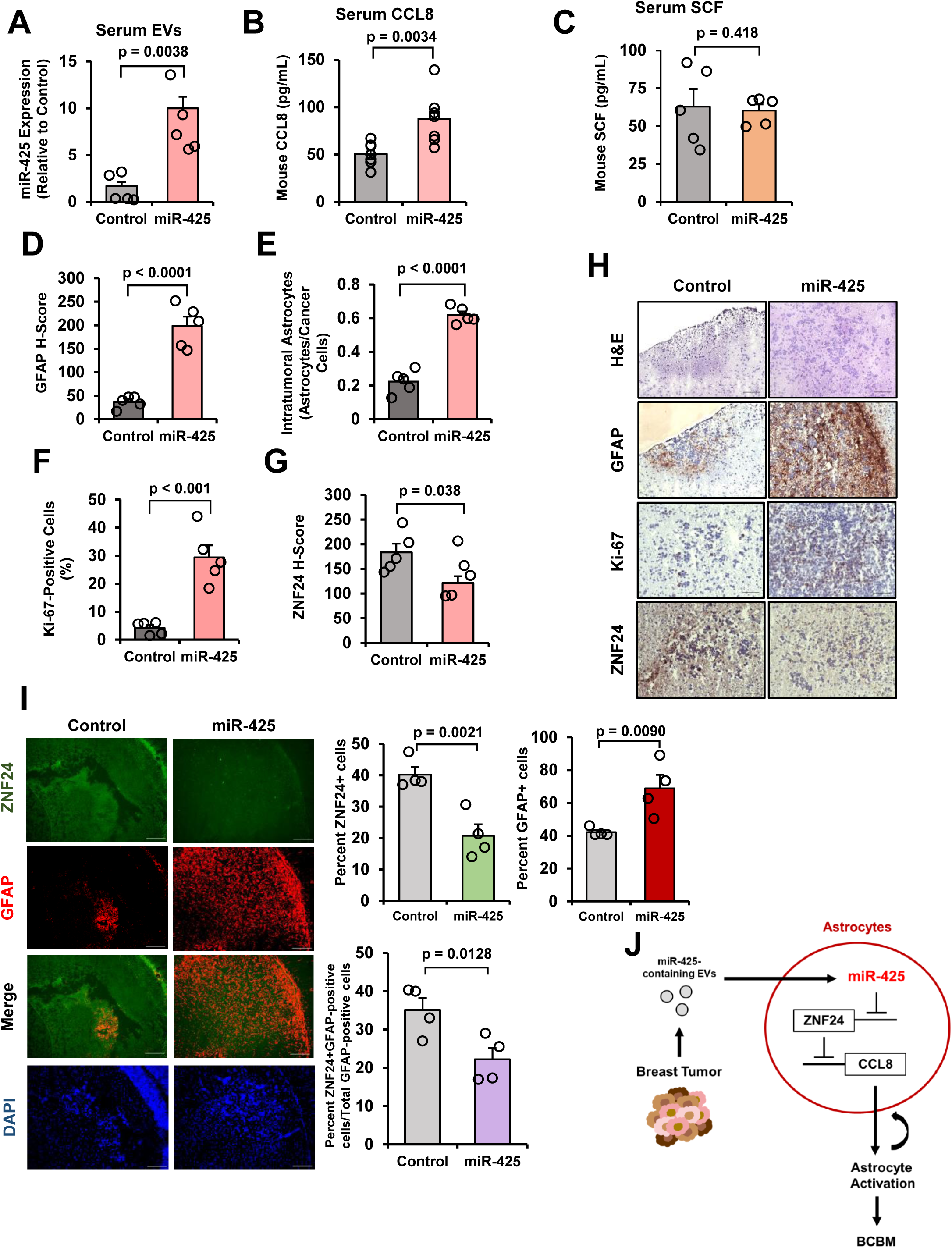
The miR-425-ZNF24-CCL8 signaling pathway is upregulated in brain metastases and activated astrocytes of mice intracardially injected with breast cancer cells overexpressing miR-425. **A)** Increased EV-miR-425 expression in mouse serum from the SKBR3-Luc-miR-425 group compared to the control group. miR-425 expression measured by RT-qPCR (N = 5 per group). **B)** SKBR3-Luc-miR-425 group mouse serum has significantly higher levels of mouse CCL8 as measured by ELISA (N = 7 per group). **C)** No significant difference in mouse SCF levels in SKBR3-Luc-miR-425 or control mice serum as measured by ELISA (N = 5 per group). **D-E)** GFAP H-score and intratumoral astrocytes are significantly higher in brain metastases from the SKBR3-Luc-miR-425 mouse group than the control in mouse brain sections as measured by IHC (N = 5 per group). **F)** Brain metastases from the SKBR3-Luc-miR-425 group have significantly increased Ki-67 positive cells as measured by IHC (N = 5 per group). **G)** Tumor-adjacent and infiltrative astrocytes in brain metastases from the SKBR3-Luc-miR-425 group have decreased ZNF24 staining measured by IHC (N = 5 per group). **H)** Representative IHC images at 20x magnification. Scale bar indicates 100 µm. **I)** Co-staining IF of ZNF24 and GFAP in mouse brain sections containing brain metastases. Representative images at 20x magnification. Scale bar indicates 100 µm. **J)** Schematic of described mechanism by which breast cancer-derived EV-miR-425 activates astrocytes by suppressing ZNF24, increasing CCL8, and thereby promoting BCBM. Fold change was calculated in Panel A. Student’s *t*-test used in Panels A-G, and I.

## DISCUSSION

In the present study, we made the following novel observations: 1) miR-425 is upregulated in serum of BCBM patients and in brain metastases, 2) high miR-425 expression and activity correlates with poor patient prognoses, 3) miR-425 activates astrocytes and miR-425-activated astrocytes promote mammospheres, 4) miR-425-activated astrocytes secrete CCL8 to further activate astrocytes and promote mammospheres, 5) miR-425 increases CCL8 secretion by directly suppressing transcription factor ZNF24, a CCL8 repressor, and 6) breast cancer-derived miR-425 promotes growth of BCBM *in vivo*.

To date, multiple miRNAs have been reported to facilitate or suppress BCBM including miR-7 [47], miR-19a [46], miR-105 [48], miR-181c [19], miR-503 [49], miR-122 [50], miR-126 [51], miR-141 [52], miR-211 [53], and miR-1290 [17]. We previously reported that breast cancer EV-derived miR-1290, which is upregulated by BCBM-promoting transcription factor tGLI1, promotes the progression of brain metastases by priming astrocytes in the brain [17]. However, no prior studies have reported a role for miR-425 in BCBM or brain metastases from any type of cancer.

miR-425 plays diverse roles in a multitude of cancers [54] including glioma [55], prostate cancer [56], acute myeloid leukemia [57], osteosarcoma [58], non-small cell lung cancer (NSCLC) [59, 60], gastric cancer [61], colorectal cancer [62], melanoma [63], and breast cancer [64, 65]. In breast cancer, miR-425 associates with poor prognosis [20], targets tumor suppressor PTEN [20, 21], promotes breast cancer cell proliferation [21], invasion *in vitro* [24], and metastasis *in vivo* [24]. In addition, EV-miR-425 is a biomarker for metastatic prostate cancer [66], breast cancer [67, 68], and colorectal cancer [23].

Interestingly, miR-425 deficiency promotes dopaminergic neuron degeneration in the brain, which is associated with Parkinson’s disease [69]. Furthermore, loss of miR-425 correlates with amyloid plaque-associated microenvironment in the brain and miR-425-deficient transgenic mice had increased neuroinflammation, neuron loss, and cognitive impairment, which is characteristic of Alzheimer’s disease [70]. These reports suggest that miR-425 plays an important role in the normal, healthy brain microenvironment, but can become dysregulated and oncogenic when derived from cancer cells.

We identified ZNF24 as a direct target of miR-425 in astrocytes. Previous studies demonstrate that ZNF24 directly binds the VEGF promoter and suppresses VEGF expression in human breast cancer cells and human microvascular endothelial cells [45, 71]. Furthermore, human breast cancer tissues with decreased ZNF24 levels compared to normal adjacent tissues had increased tumor angiogenesis [45]. ZNF24’s tumor suppressive role has also been reported in gastric cancer [72]. In contrast, ZNF24 facilitates EMT in prostate cancer [73], suggesting that ZNF24 functions may be tumor-or tissue-specific.

In this study, we found that CCL8 (or MCP-2) is highly secreted by miR-425-activated astrocytes and the significant increase is due to miR-425 suppression of ZNF24, which directly suppresses CCL8 expression. Previous studies demonstrate that CCL8 is upregulated in breast cancer tissues compared to normal tissues [74], and contributes to breast cancer migration [75], invasion [75], and metastasis [75]. In the brain, CCL8 is secreted by tumor-associated macrophages to promote glioblastoma growth [76] and CCL8 is a target for miR-146a in HIV-1-infected human microglial cells [77]. There are currently no reports of ZNF24 directly repressing CCL8 expression in any cancer type.

Our results indicate that breast cancer cells overexpressing miR-425 promote larger brain metastases compared to the control group. Interestingly, we did not observe a significant difference in the frequency of brain metastases between both groups (data not shown), suggesting that breast cancer-derived miR-425 promotes brain metastasis growth after colonization and may not directly impact breast cancer colonization in the brain. Our study identifies, for the first time, the important brain-tropic role of miR-425 in breast cancer.

Since the discovery of the first miRNA in 1993 [78], the field of miRNAs in human cancer has flourished. There are over 2000 mature miRNAs identified in the human body with many being studied in human cancers and other diseases [15]. Though there are numerous miRNA-based therapeutic clinical trials, currently no miRNA therapeutic has been FDA-approved, suggesting that we still have much to learn about the roles of miRNAs in cancer [79]. Our study aimed to address mechanisms that drive BCBM and found that EV-miR-425, which is upregulated in BCBM patient serum, can activate astrocytes in the brain to promote the progression of brain metastases. Our results indicate that miR-425 is an important therapeutic target for BCBM that should be examined further in brain metastases and potentially other brain cancers. In summary, our novel findings provide new knowledge of miR-425 functions, and mechanisms by which breast cancer activates astrocytes to facilitate brain metastasis.

## MATERIALS AND METHODS

### Cell lines, lentiviruses, miRNA mimics, and expression plasmids

SKBR3 cells were purchased from the American Type Culture Collection (ATCC). CN34 and CN34-BRM cells, the brain metastatic variant derived from TNBC CN34 cells, were a kind gift from Dr. Joan Massagué at the Sloan Kettering Institute [26]. SKBRM cells, the brain metastatic variant derived from HER2-enriched SKBR3 cells, were previously established by Drs. Fei Xing and Kounosuke Watabe [80]. The E6/E7/hTERT immortalized human astrocytes were a kind gift from Dr. Russell O. Pieper at the University of California, San Francisco. Cells were transfected using mirVana mimics and the corresponding inhibitors were purchased from ThermoFisher Scientific (Waltham, MA). The miRNA mimics include: negative control (4464058), hsa-miR-107 (4464066, MC10056), and hsa-miR-425-5p (4464066, MC11575). The miRNA inhibitors include: negative control (4464076), hsa-miR-107 (4464084, MH10056), and hsa-miR-425-5p (4464084, MH11575). Additional plasmids used for cell transfection include: ZNF24 overexpression and ZNF24 shRNA plasmids. ZNF24 overexpression and control plasmids were purchased from VectorBuilder (Chicago, IL) and custom designed to overexpress human ZNF24 (NM_006965.4; E. coli (VB221017-1240ueu)) or control (ORF_Stuffer; E. coli (VB900122-0478rbb)). ZNF24 shRNA (scbt; sc-76969-SH) and control (scbt; sc-108060) plasmids were purchased from Santa Cruz Biotechnology (Dallas, TX). Breast cancer cell lines stably expressing hsa-miR-425 were generated using the human pre-microRNA expression construct control and lenti-miR-425 purchased from System Biosciences (PMIRH000PA-1, PMIRH425-PA-1; Palo Alto, CA).

Lentiviruses were generated by GenScript (Piscataway, NJ) and then transduced into SKBR3 cells stably expressing luciferase. Cells were sorted using the GFP reporter. All cell lines were authenticated using standard methods, tested using the MycoStrip Mycoplasma Detection Kit (InvivoGen; San Diego, CA), treated if necessary (10-799-050-001; Sigma-Aldrich; St. Louis, MO), and tested again prior to use.

### Extracellular vesicle (EV) isolation and nanoparticle tracking analysis (NTA)

EV-free FBS preparation was carried out as previously described [17]. For cells, EVs were isolated using the ExoQuick-TC ULTRA Kit following the manufacturer’s instructions (EQULTRA-20TC-1; System Biosciences). For patient or mouse serum, EVs were isolated using the ExoQuick ULTRA Kit following the manufacturer’s instructions (EQULTRA-20A-1; System Biosciences). Particle size distribution was measured by NTA using the Nanosight NS50 (Malvern Instruments, UK).

### Reverse transcription quantitative real-time PCR (RT-qPCR)

RT-qPCR was performed as previously described [17]. RNA was isolated using the RNeasy Mini Kit (Qiagen; 74106), cDNA was synthesized using SuperScript III Reverse Transcriptase (Invitrogen; 18080093), and qPCR was performed using GoTaq qPCR Master Mix (Promega; A6001). Relative quantification was performed using the 2^-ΔΔCT^ method. qPCR primers utilized are listed in Supplementary Table I.

### MiRNA isolation and RT-qPCR

MiRNAs were isolated using the miRNeasy Mini Kit (Qiagen; 217004), as previously described [17]. Primer sets were purchased from ThermoFisher (A25576): hsa-miR-107 (478254_mir), hsa-miR-425-5p (478094_mir), hsa-miR-451a (478107_mir), and hsa-miR-361-5p (478056_mir).

hsa-miR-451a (478107_mir), and hsa-miR-361-5p (478056_mir) were used as internal controls, and 2^-ΔΔCT^ method was used for relative quantification. qPCR was carried out on the Applied Biosystems QuantStudio 3 Real-Time PCR System (ThermoFisher; A28137).

### MiRNA overexpression and inhibition

Cells were transduced using miRNA mimics or inhibitors (described above) and as previously described [17]. Briefly, cells were seeded at 1-3 x 10^5^ cells in 6-well culture plates and incubated overnight at 37°C. Next, the Lipofectamine RNAiMAX Transfection Reagent (13778150) and the miRNA mimic/inhibitor were mixed with Opti-MEM separately and left to equilibrate for 5 minutes. Both tubes were mixed, incubated for 15 minutes, and then added dropwise to the 6-well plates containing 1750 µL of fresh media with cells. After 5-6 hours, the transfection mix was replaced with fresh cell culture media, and cells were incubated for another 42 hours before harvesting.

### Immunoblotting and Immunohistochemistry (IHC)

Immunoblotting was performed as previously described [81]. Lysates were subjected to 10% or 4-20% precast sodium dodecyl sulfate-polyacrylamide gel (Bio Rad; 5671033 or 5671093) depending on the size of each protein. Antibodies used for immunoblotting include: CCL8/MCP-2 (Abcam; ab155966; 1:500), SCF (Cell Signaling; CST2093S; 1:1000), Vinculin (Cell Signaling; CST13901S; 1:10,000), and ZNF24 (Santa Cruz Biotechnology; sc-392259; 1:500). IHC was conducted as previously described [82, 83]. IHC antibodies include Ki-67 (CST; 9027S), GFAP (abcam; 68428), ZNF24 (Proteintech; 11219-1-AP), and PTEN (Cell Signaling; CST9559). Histochemical scores (H-scores) were calculated using the following equation: H-score = A x B, where A represents percent positivity (A%, A = 1-100) and B represents intensity (B = 0-3).

### Breast cancer mammosphere assays

Mammosphere assays were performed as previously described [17]. Breast cancer cells were transfected with miRNA mimics or inhibitors (described above), then collected, counted and seeded in 24-well ultra-low attachment plates (Corning) at 500-2000 cells per well. Astrocyte-conditioned media (CM) was obtained from cells cultured in serum-free medium for 24-48 hours.

### Glial fibrillary acidic protein (GFAP) and ZNF24 immunofluorescence (IF)

GFAP immunofluorescence for human astrocytes was carried out as previously described [17]. GFAP antibody (Cell Signaling; 3656) was used at a 1:25 dilution and DAPI antibody at 1:1000 dilution. ZNF24 immunofluorescent antibody (Bioss; bs18508R-FITC) was used at a 1:50 dilution, GFAP antibody (Cell Signaling; 3656) was used at 1:50 dilution, and DAPI antibody at 1:1000 dilution on 10 µm sections of mouse brains with brain metastases. Mouse brain tissues were fixed in acetone, washed with PBS-T (0.3%), incubated with mouse-mouse blocking reagent (ScyTek; MTM015) for 30 minutes, incubated with blocking (5% goat serum/PBS-T) for 1 hour, and incubated with primary antibodies (see above) overnight at 4°C. The next day, slides were washed with PBS, incubated with DAPI, washed again and then mounted using aqueous mounting medium. All slides were imaged at 4x or 20x using the ImageXpress Pico or Keyence BZ-X810 microscopes.

### Cytokine arrays

Human astrocytes were transfected with control or miR-425 mimic (described above). After 48 hours, cells were washed with PBS, then serum-free media was replaced and incubated for 24 hours. CM was collected, centrifuged at 1250 rpm, supernatant was transferred, and 1 mL aliquots were stored at −80°C. CM was used in the Human Cytokine Antibody Array (Abcam; ab133998) following the manufacturer’s instructions.

### Enzyme-linked immunosorbent assays (ELISA)

Human astrocytes were transfected with control or miR-425 mimic (described above) for 48 hours. Cells were washed with PBS, incubated with serum-free media for 24 hours, CM was collected, centrifuged at 1250 rpm, and supernatant was transferred and stored at −80°C in 500 µl aliquots. CM was used in the Human MCP2 ELISA Kit (abcam; ab223856) and Human SCF ELISA Kit (abcam; ab176109) following the manufacturer’s instructions. For mouse serum, mouse whole blood was collected at euthanization, centrifuged at 2000 x g for 15 mins at 4°C, serum was transferred to new tubes and stored for EV isolation and ELISAs. Serum was used in the Mouse MCP2 ELISA Kit (CCL8) (abcam; ab203366) and Mouse SCF ELISA Kit (abcam; ab197750) following the manufacturer’s instructions.

### Chromatin immunoprecipitation-qPCR (ChIP-qPCR) assays

ChIP-qPCR assays were performed as previously described using the EZ-ChIP Kit (Millipore Sigma; 17-371) [17, 84]. Astrocytes were transfected with control or ZNF24 overexpression plasmids (VectorBuilder; VB221017-1240ueu) and CCL8 promoter (GeneCopoeia; HPRM46246-PG02) using X-tremeGENE HP DNA Transfection Reagent (Millipore Sigma; 6366244001). Control and ZNF24 transfected cell lysates were immunoprecipitated using the ZNF24 antibody (scbt; sc-393359), or the normal mouse IgG provided in the EZ-ChIP Kit. Primer sequences amplifying the human CCL8 promoter in two different regions (−69 and -205 bases upstream of the transcription start site) include 5’ – CCCACTGTGTGTGAACCAAG – 3’ (Region 1 forward), 5’ – GCACATGCTGGTCTTGTAGC – 3’ (Region 1 reverse), 5’ – TAAAGGAGCTTGCCTGGCTT – 3’ (Region 2 forward), and 5’ – CCTAAGGCTTTGCTGGTGCT – 3’ (Region 2 reverse). Mouse normal IgG served as negative immunoprecipitation controls. Input chromatins were used as controls for qPCR following the ChIP assay.

### Luciferase reporter assays

Dual luciferase reporter assays were carried out as previously described [17, 83]. Astrocytes were transfected with control or miR-425 mimic, control vector or ZNF24 3’UTR (GeneCopoeia; Cs-HmiT119980-MT05-01), control vector or ZNF24 overexpression (VB221017-1240ueu), and control promoter or CCL8 promoter (GeneCopoeia; HPRM46246-PG02) constructs using X-tremeGENE HP DNA Transfection Reagent (Millipore Sigma; 6366244001). The Secrete-Pair Dual Luminescence Assay Kit (GeneCopoeia; LF032) was carried out following the manufacturer’s instructions. Relative 3’UTR or promoter activity was determined by normalizing the luciferase activity to secreted alkaline phosphatase (SEAP) and the respective transfection control.

### MiRNA expression profiling, gene expression profiling, and metastasis-free survival analysis

MiRNA expression datasets were obtained from Gene Expression Omnibus (GEO): GSE68373, GSE37407, GSE134108, GSE73002, GSE40525, and GSE22220. GSE12276/2034/2603/5327/14020 breast cancer patient datasets were compiled into one GEO dataset as previously described [26].

### Intracardiac inoculation mouse model

All animal experiments were conducted as approved by the UTHealth Houston Center for Laboratory Animal Medicine and Care (CLAMC). Isogenic SKBR3 lines with vector or miR-425 overexpression (SKBR3-Luc-miR-Control/425) were injected (2×10^5^ cells) into the left cardiac ventricle of female athymic mice and monitored using an *In Vivo* Imaging System (IVIS). Brain and bone metastasis occurrence and growth was monitored through twice-weekly IVIS imaging and mice were euthanized at humane endpoint or end of study. Blood serum was collected, all organs imaged *ex vivo*, and mouse brains were stored in optimal cutting temperature (O.C.T) compound for further analysis.

### Statistical analyses

Log-Rank (Mantle-Cox) test, one-way ANOVA, two-way ANOVA, and student’s *t*-test were performed using GraphPad Prism 9 as previously described [83]. N = 3 experimental replicates unless otherwise indicated. Results are represented as ± SE.

## AUTHOR CONTRIBUTIONS

**Grace L. Wong**: conceptualization, data curation, formal analysis, methodology, writing – original draft, writing – review & editing, and visualization. **Munazza S. Khan**: writing – review & editing, and visualization. **Sara Manore**: animal handling. **Shivani Bindal**: animal handling. **Ravi Singh**: methodology, data curation, and formal analysis. **Hui-Wen Lo**: conceptualization, funding acquisition, investigation, methodology, project administration, resources, supervision, validation, visualization, writing – review & editing. All authors have read and agreed to the published version of the manuscript.

## Supporting information

Original Western Blots

## ACKNOWLEDGMENTS

The authors would like to acknowledge Drs. Fei Xing and Kounosuke Watabe for the SKBRM cells, Dr. Joan Massagué for the CN34-BRM cells, and Dr. Russell O. Pieper for the immortalized human astrocytes. We acknowledge funding support for this study from NIH grants R01CA228137 (H-WL) and R21CA286225 (H-WL), DoD grants W81XWH-19-1-0072 (H-WL), W81XWH-20-1-0044 (H-WL), W81XWH-19-1-0753 (H-WL), and HT9425-24-1-0889 (H-WL), as well as, MetaVivor Translational Research Grants (H-WL) and UTStars (H-WL).

