## Supplemental Figures and Table for "Extracellular vesicle-derived miR-425-5p (miR-425) activates astrocytes in the brain to promote breast cancer brain metastasis via the novel miR-425-ZNF24-CCL8 signaling axis"

### Supplementary Figures

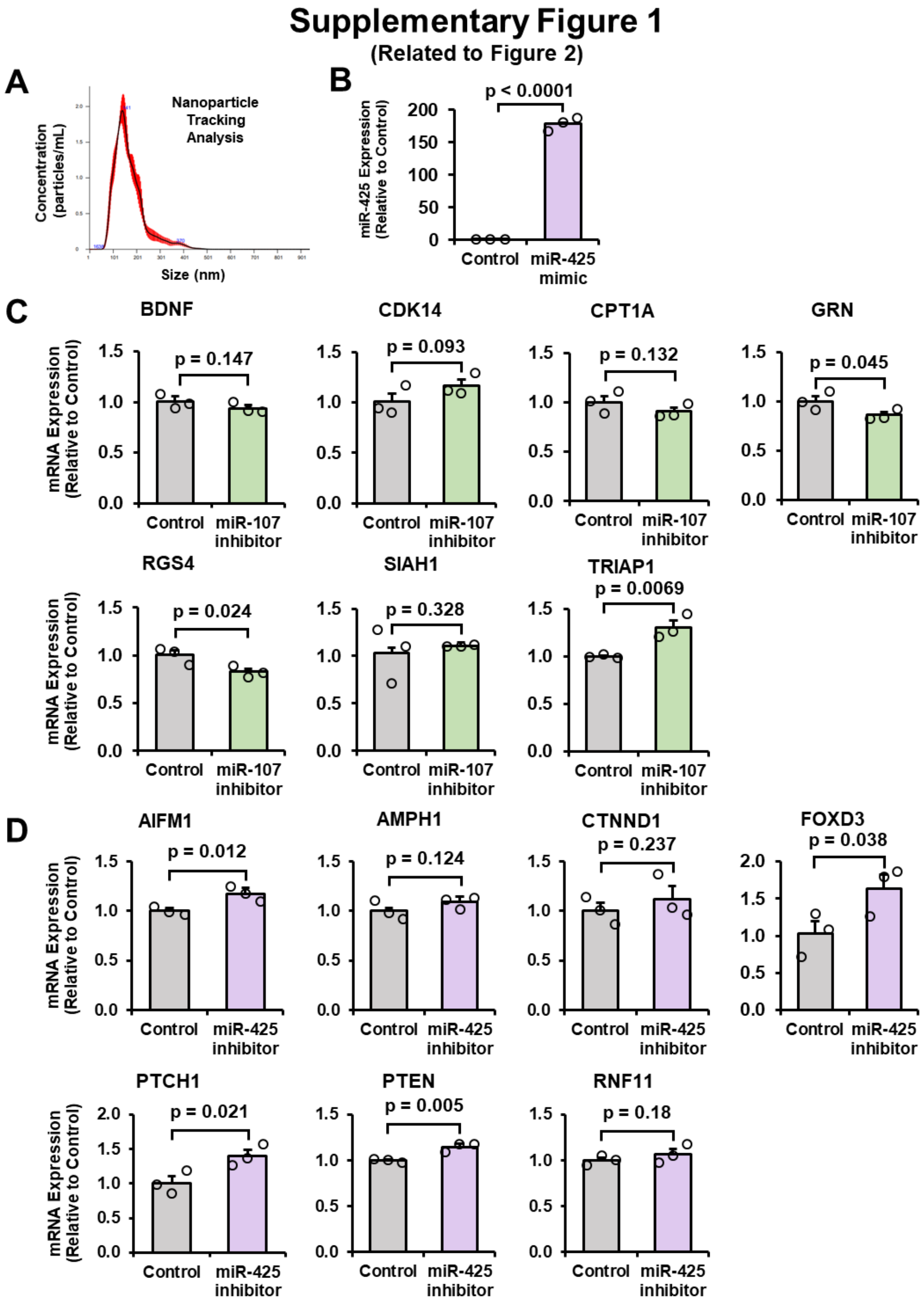


**Supplementary Figure 1: Validation of EV isolation, miR-425 overexpression, and miR-107 and miR-425 inhibitors. A)** EV isolation via the ExoQuick Kit was validated via Nanoparticle Tracking Analysis (NTA). **B)** Validation of miR-425 overexpression in CN34 cells via miR-RT-qPCR. **C)** miR-107 inhibitor was validated via RT-qPCR of previously published miR-107 target genes. **D)** miR-425 inhibitor was validated via RT-qPCR of previously published miR-425 target genes. Fold change was calculated in Panels B-D. Student's *t*-test was used in Panels B-D. N = 3 experimental replicates unless otherwise indicated.

### Supplementary Figure 2

(Related to Figure 4)

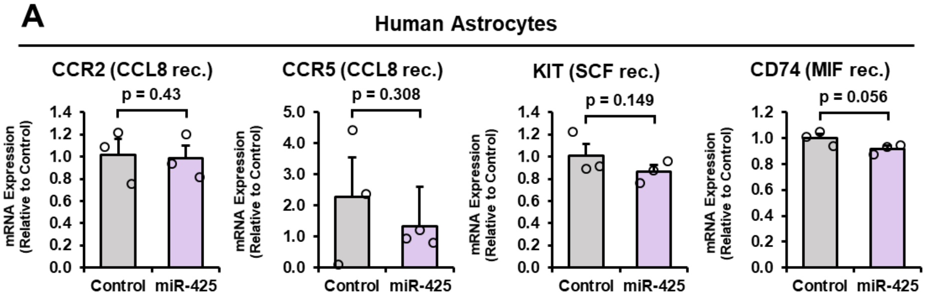


**Supplementary Figure 2: CCL8, SCF, and MIF receptor expression in human astrocytes. A)** CCR2 (CCL8 receptor), CCR5 (CCL8 receptor), KIT (SCF receptor), and CD74 (MIF receptor) mRNA expression in astrocytes transfected with the miR-425 mimic. mRNA levels measured with RT-qPCR. Fold change was calculated in Panel A. Student's *t*-test was used in Panel A. N = 3 experimental replicates unless otherwise indicated.

### Supplementary Figure 3

(Related to Figure 8)

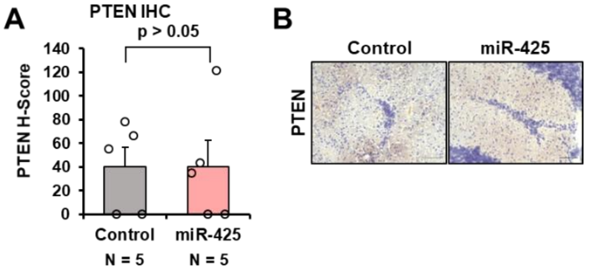


**Supplementary Figure 3: PTEN protein expression in brain metastases from mice intracardially injected with breast cancer cells overexpressing miR-425. A)** PTEN expression is not significantly different in brain metastases from the control or miR-425-overexpressing groups. **B)** Representative IHC images at 20x magnification. Scale bar indicates 100 µm.

### Supplementary Tables

#### Supplementary Table I:

#### RT-qPCR Primer Sequences:

| Target Gene | Forward Sequence (5'-3') | Reverse Sequence (5'-3') |
|---|---|---|
| AIFM1 | TTGAGAATGGTGGTGTGGCT | AGACTTCTTGGAGTACCTCCTGT |
| AMPH1 | CGAGAACTCCGAGGATATTTAGC | CCCATACCAGTCAGGCTCAT |
| BCOR | CGCTCCTCGCTGAACGC | CGCCATGTTGACGGTTCGC |
| BDNF | AGATCTTGGGGGAAACACTGC | TAGGGCTTTCTTTCACCGGG |
| CCL8 | TGTCCCAAGGAAGCTGTGAT | TGGAATCCCTGACCCATCTCT |
| CDK14 | GATGTGTGACCTCATTGAGCC | CAATGCGACTGAAACTCTCCG |
| CPT1A | TGTCCAGCCAGACGAAGAAC | ATCTTGCCGTGCTCAGTGAA |
| CTNND1 | ATGAGTGGTTCTCCAGAGGGA | GCAGAGCAGAGCGGATGTAT |
| CREB1 | GTGACGGAGGAGCTTGTACC | GGACTTGAACTGTCTGCCCA |
| FOXD3 | GCAACTACTGGACCCTGGAC | CTGTAAGCGCCGAAGCTCT |
| GAPDH | ACTGCCAACGTGTCAGTGG | GTGTCGCTGTTGAAGTCAGA |
| GFAP | CTGCTGCCTTTAGTCGCTGA | CTGCGGGTGGAATTTGGTGA |
| GRN | ATCTTTACCGTCTCAGGGACTT | CCATCGACCATAACACAGCAC |
| KITLG | AGCCAGCTCCCTTAGGAATGA | TGCCCTTGTAAGACTTGGCTG |
| MIF | ATCGTAAACACCAACGTGCC | TGCTGTAGGAGCGGTTCTG |
| PTCH1 | GGGTGGCACAGTCAAGAACAG | TACCCCTTGAAGTGCTCGTACA |
| PTEN | AACTTGCAATCCTCAGTTTG | CTACTTTGATATCACCACACAC |
| RGS4 | AGAGGAAAGGCATTGGGAGTC | GCTAAGCCTGTAGGGGTCTC |
| RNF11 | TCTCCCTGCTTCACGAGTCT | AGTCTGGCTAGGTGTTGGGT |
| SIAH | AGGACCCTACCCAGTGAATCT | GACCCAAATTCGCGTCTGAG |
| TRIAP1 | ATTGCCAGCTCTCAACCCAA | AAGGCAAATGAAGCGAGCAC |
| VEGFA | TGTCTAATGCCCTGGAGCCT | TTAACTCAAGCTGCCTCGCC |
| ZNF24 | GGAGGTTTGCGCCGGAGT | ACAGACGGCAAAGTTCTCGG |
