## Supplementary figures and images for "Extracellular vesicle-derived miR-425-5p (miR-425) activates astrocytes in the brain to promote breast cancer brain metastasis via the novel miR-425-ZNF24-CCL8 signaling axis"

### Original Western Blots

#### ORIGINAL WESTERN BLOTS

##### Figure 5 C

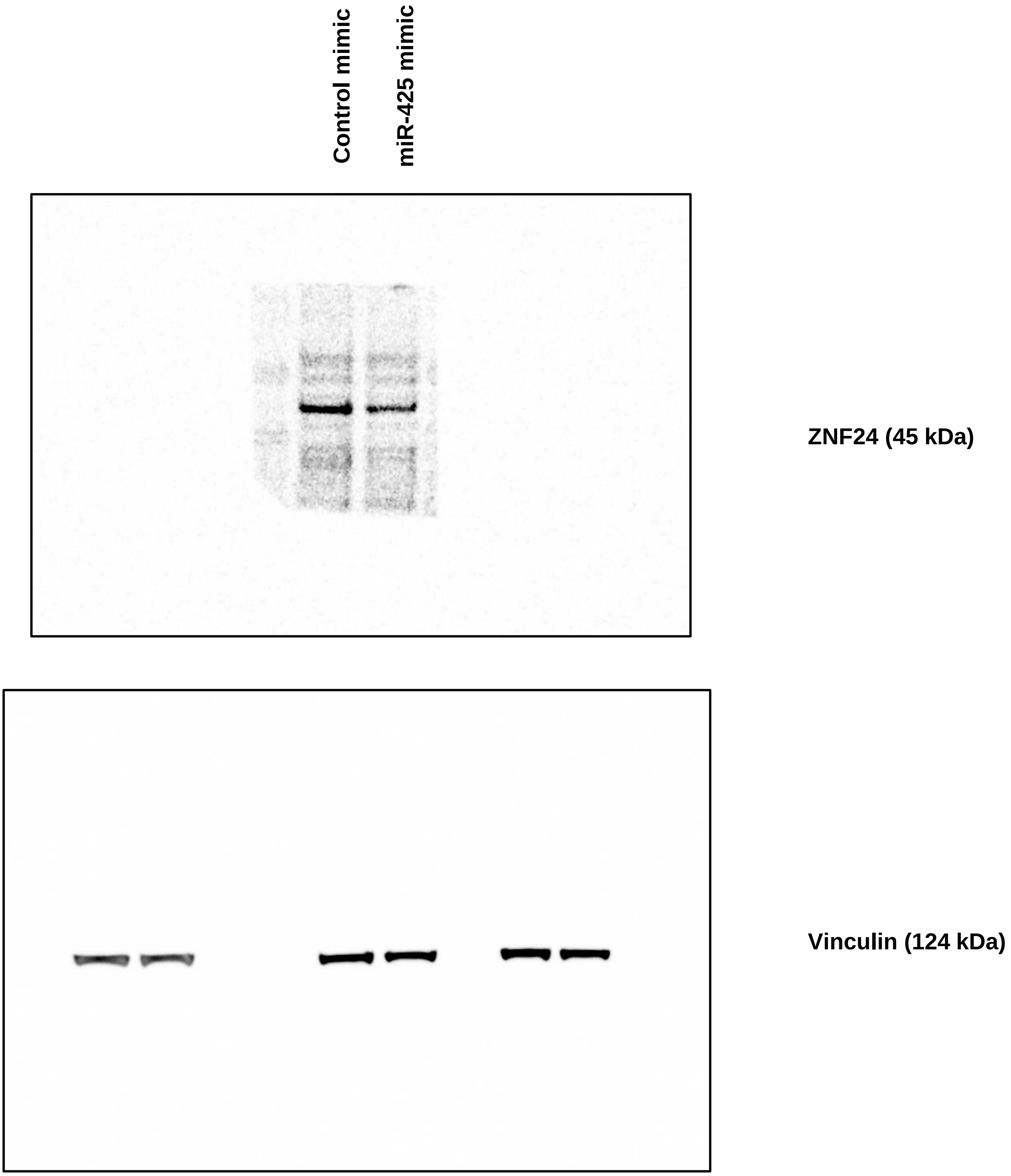

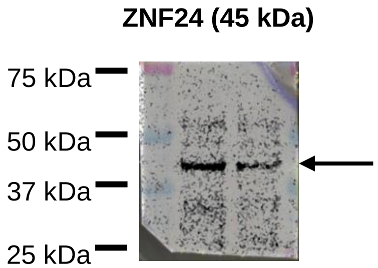

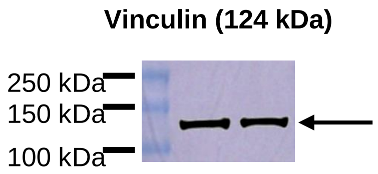

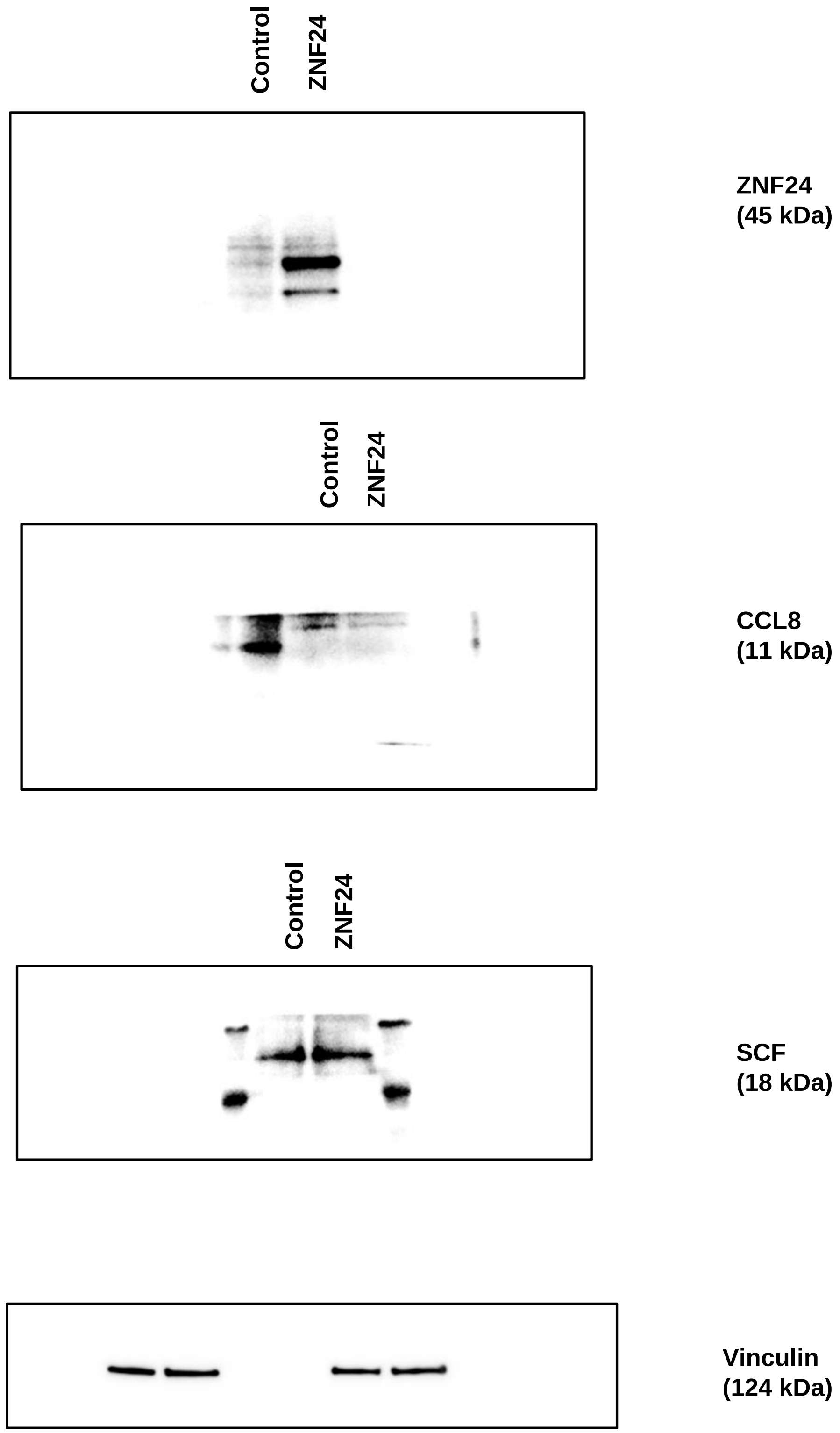

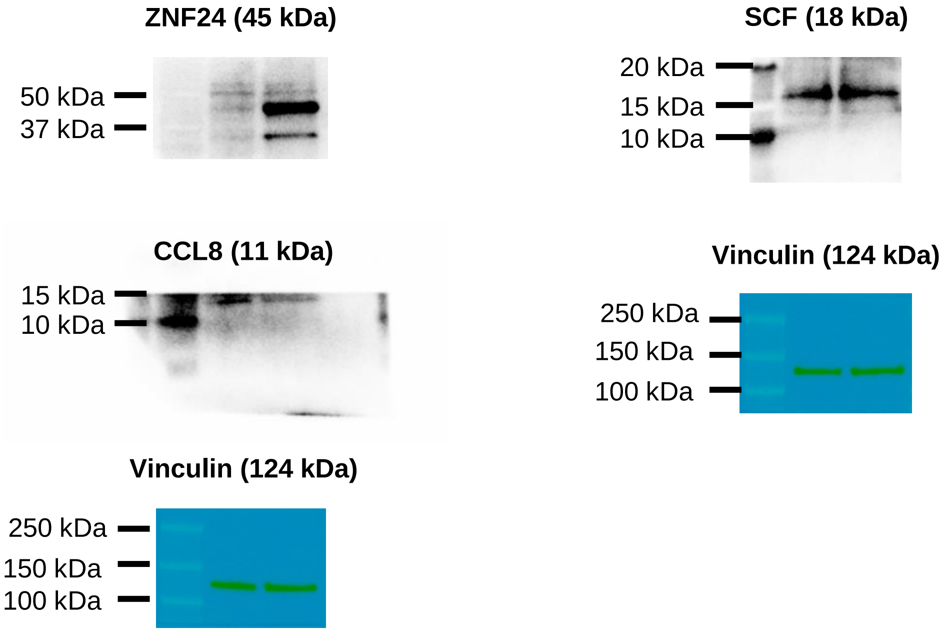

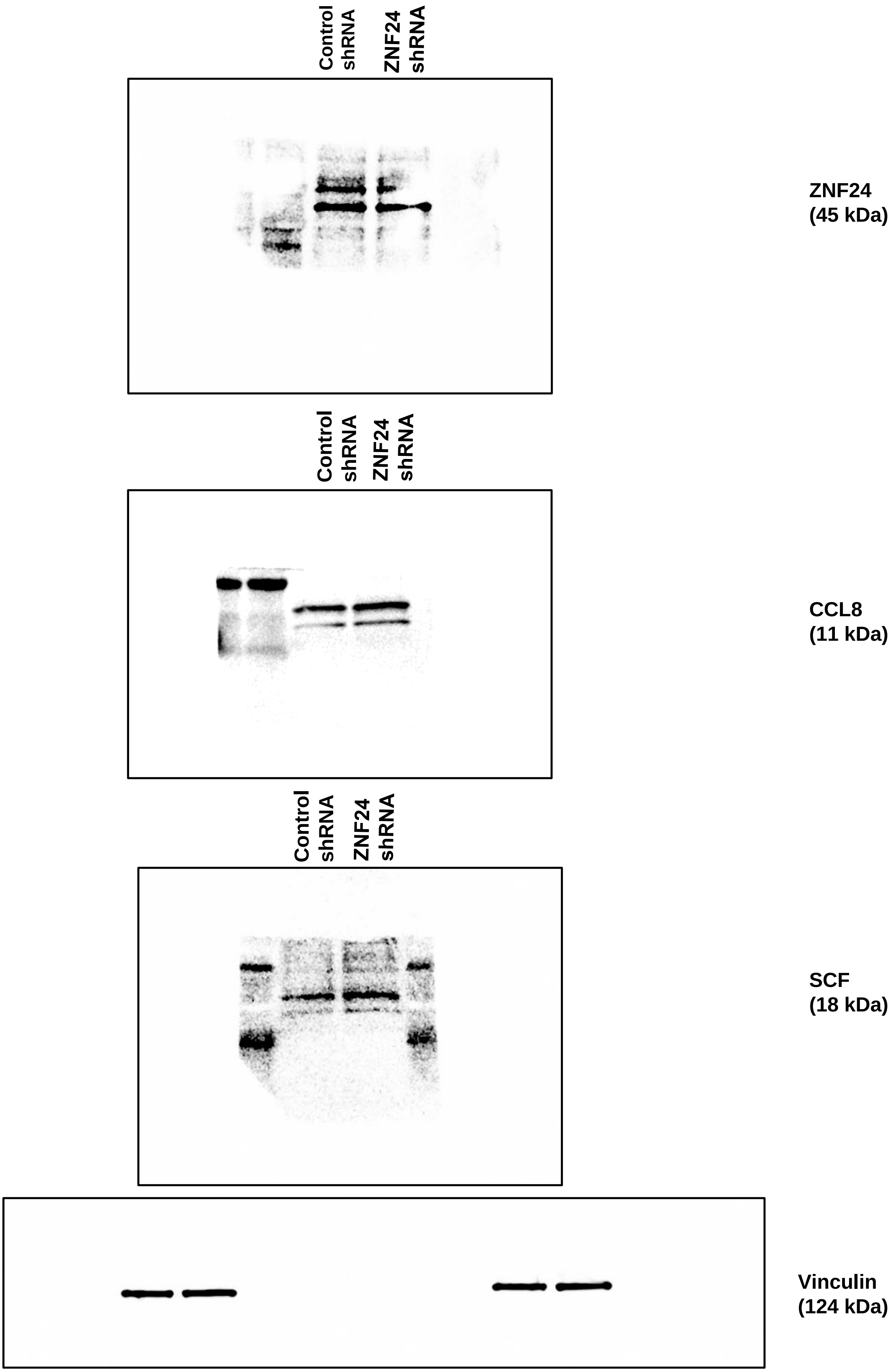

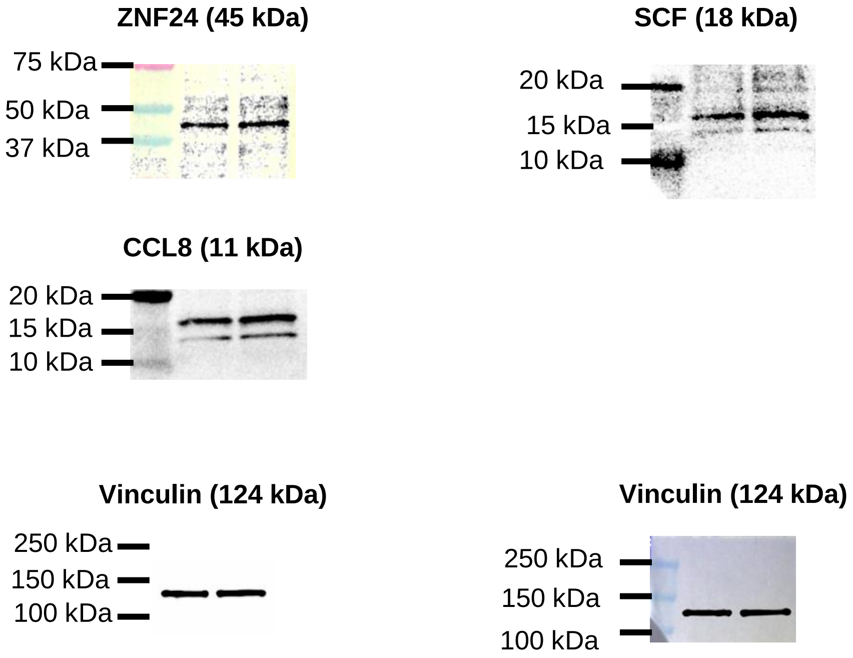
